# Pharmacological inhibition of soluble epoxide hydrolase as a new therapy for Alzheimer’s Disease

**DOI:** 10.1101/605055

**Authors:** Christian Griñán-Ferré, Sandra Codony, Eugènia Pujol, Jun Yang, Rosana Leiva, Carmen Escolano, Dolors Puigoriol-Illamola, Júlia Companys-Alemany, Rubén Corpas, Coral Sanfeliu, M. Isabel Loza, José Brea, Christophe Morisseau, Bruce D. Hammock, Santiago Vázquez, Mercè Pallàs, Carles Galdeano

**Affiliations:** Pharmacology Section. Department of Pharmacology, Toxicology and Medicinal Chemistry, Faculty of Pharmacy and Food Sciences, and Institut de Neurociències, University of Barcelona, Av. Joan XXIII, 27-31, E-08028 Barcelona, Spain; Laboratori de Química Farmacèutica (Unitat Associada al CSIC), Department de Farmacologia, Toxicologia i Química Terapèutica, Facultat de Farmàcia i Ciències de l’Alimentació, and Institute of Biomedicine (IBUB), Universitat de Barcelona, Av. Joan XXIII, 27-31, E-08028 Barcelona, Spain; Department of Entomology and Nematology and Comprehensive Cancer Center, University of California, Davis, California, USA; Institute of Biomedical Research of Barcelona (IBB), CSIC and IDIBAPS, Barcelona, Spain; CIBER Epidemiology and Public Health (CIBERESP), Madrid, Spain; Innopharma screening platform. Biofarma research group. Centro de Investigación en Medicina Molecular y Enfermedades Crónicas (CIMUS). Universidad de Santiago de Compostela. Spain; Department of Pharmacy and Pharmaceutical Technology and Physical Chemistry, Faculty of Pharmacy and Food Sciences and Institute of Biomedicine (IBUB), University of Barcelona, Av. Joan XXIII, 27-31, E-08028 Barcelona, Spain

## Abstract

The inhibition of the enzyme soluble epoxide hydrolase (sEH) has demonstrated clinical therapeutic effects in several peripheral inflammatory-related diseases, with two compounds that have entered clinical trials. However, the role of this enzyme in the neuroinflammation process has been largely neglected. Herein, we disclose the pharmacological validation of sEH as a novel target for the treatment of Alzheimer’s Disease (AD). Of interest, we have found that sEH is upregulated in brains from AD patients. We have evaluated the cognitive impairment and the pathological hallmarks in two models of age-related cognitive decline and AD using three structurally different and potent sEH inhibitors as chemical probes. Our findings supported our expectations on the beneficial effects of central sEH inhibition, regarding of reducing cognitive impairment, tau hyperphosphorylation pathology and the number of amyloid plaques. Interestingly, our results suggest that reduction of inflammation in the brain is a relevant therapeutic strategy for all stages of AD.

## INTRODUCTION

Chronic inflammation is recognized as a key player in both onset and progression of Alzheimer’s Disease (AD) (*1-3*). Indeed, 16% of the investment in ongoing clinical trials for AD is related to inflammation (*4*). Neuroinflammation is intimately linked to the oxidative stress (OS) associated with AD (*5,6*), controlling the interactions between the immune system and the nervous system (*7,8*). However, several antioxidant therapies and non-steroidal anti-inflammatory drugs have failed in clinical trials. Therefore, it is of vital importance to expand the scope towards novel targets, preferably related with several pathophysiological pathways of the disease (*9*).

Epoxyeicosatrienoic acids (EETs) mediate vasodilatation, reduce inflammation, attenuate OS and block the pathological endoplasmic reticulum (ER) stress response (*10,11*). The soluble epoxide hydrolase enzyme (sEH, EC 3.3.2.10, EPHX2), widely expressed in relatively high abundance in the murine and human brains (*12,13*), converts EETs and other epoxyfatty acids (EpFA) to their corresponding dihydroxyeicosatrienoic acids (DHETs), whereby diminishing, eliminating, or altering the beneficial effects of EETs (*14*) (Fig. 1).

**Fig. 1.**
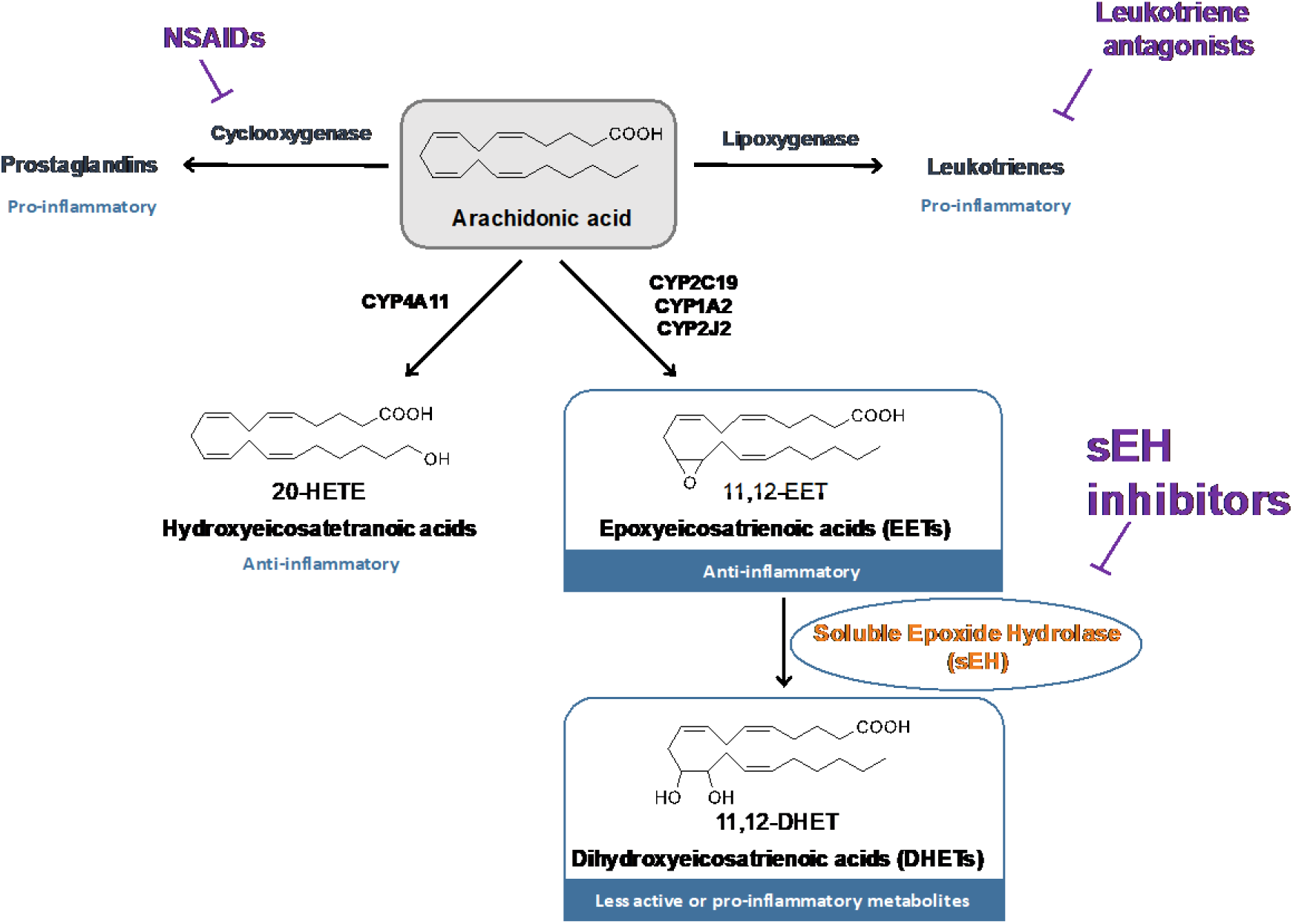
The arachidonic acid cascade. The arachidonic acid (AA) cascade is a group of metabolic pathways in which AA and other polyunsaturated fatty acids are the central molecules. Metabolism via the cyclooxygenase (COX) and lipoxygenase (LOX) pathways gives rise to largely pro-inflammatory and pro-algesic metabolites. Both pathways have been pharmaceutically targeted. CYP enzymes either hydroxylate or epoxidize AA leading to hydroxyeicosatetranoic acids (HETEs) or epoxyeicosatrienoic acids (EETs), respectively. The latters, which are endowed with potent anti-inflammatory properties, are rapidly subjected to hydrolysis to their corresponding diols by the soluble epoxide hydrolase (sEH) enzyme. Inhibitors of sEH block this degradation and stabilize EETs levels *in vivo* (*14*). Major CYPs that oxidize AA are listed in the figure, but many others make a contribution.

Considering that several lines of evidence underline a broad involvement of signaling by EETs and other EpFA in the central nervous system (CNS) function and disease (*15,16*), we hypothesized that brain penetrant sEH inhibitors would stabilize EETs in the brain, resulting in a reduction of reactive oxygen species (ROS) and diminished neuroinflammation and neurodegeneration leading to a positive outcome in AD. Of interest, we found that sEH is overexpressed in brain tissues from human AD patients in Braak stage V (Fig. 2A and table S1). We also found sEH overexpressed in the hippocampus tissues of two murine models, 5xFAD (early on-set AD, Fig. 2B) and SAMP8 (late onset AD, Fig. 2B), supporting a crucial role of this enzyme in the disease.

**Fig. 2.**
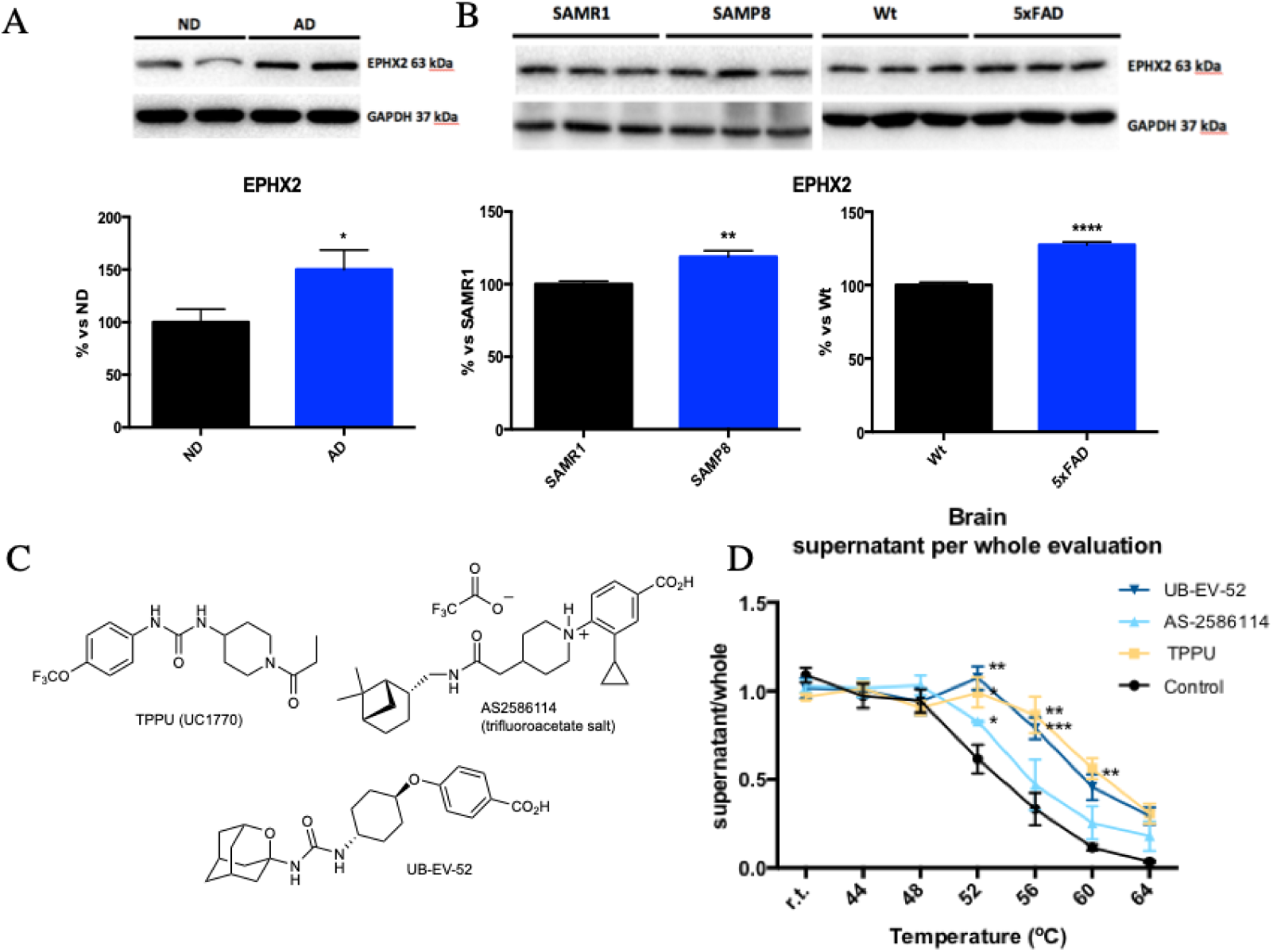
Soluble epoxide inhibition and its relevance in AD. **(A)** Immunoblot of sEH (EPHX2) of human brains from patients. Groups were compared by Student t-test (n = 3/4 (AD)). *p<0.05 vs. non-demented. **(B)** Immunoblot of sEH (EPHX2) in the hippocampus of SAMP8 mice (groups were compared by Student t-test, n = 12-14, **p<0.01 vs. SAMR1) and 5xFAD mice (groups were compared by Student t-test, n = 12-14, ****p<0.0001 vs. Wt). **(C)** Chemical structure of the sEH inhibitors employed. **(D)** CETSA experiments to monitor brain target engagement. Groups were compared by Student t-test or Two-Way ANOVA and post-hoc Dunnett’s, n = 3 per group, *p<0.05, **p<0.01 and ***p<0.001 vs. Control.

## RESULTS

### On-target drug inhibition of sEH

We evaluated the pathological hallmarks and the cognitive impairment associated with AD using three structurally different sEH inhibitors as chemical probes (*17*): TPPU (UC1770, IC_50_ for human sEH = 3.7 nM) (*18*), AS-2586114 (IC_50_ for human sEH = 0.4 nM) (*19*), and UB-EV-52 (IC_50_ for human sEH = 9 nM) (*20*) (Fig. 2C). Previous pharmacokinetic data suggest that TPPU, a very well characterized sEH inhibitor, can enter into the brain (*21-22*). It is known that AS-2586114 has a prolonged action *in vivo* and ability to cross the blood–brain barrier (*23-24*). UB-EV-52 is a new inhibitor somewhat related with previously reported adamantane-derived sEH inhibitors such as *t*-AUCB (*25*) and the clinically studied AR9281 (UC1153) (*26*). To determine whether UB-EV-52 possesses drug-like characteristics, we performed *in vitro* ADMET assays. We found that UB-EV-52 has excellent solubility (>100 µM at 37 °C in 5% DMSO: 95% PBS buffer), good microsomal stability (table S2) and does not inhibit cytochromes neither hERG (table S3). Of relevance, some cytochromes are potential *off*-target of sEH (Fig. 1), since they are situated upstream in the arachidonic acid cascade. UB-EV-52 showed less than 5% inhibition of the studied cytochromes (CYP1A2, CYP2C9, CYP2C19, CYP3A4 and CYP2D6) at 10 µM (table S3). In order to characterize the toxicity of UB-EV-52, we evaluated cell viability in human neuroblastoma SH-SY5Y cells, using a MTT assay for cell metabolic activity and a propidium iodide stain assay for cell death. In both assays, UB-EV-52 showed no cytotoxicity at 1, 10, 50 and 100 µM.

To evaluate whether sEH is the direct binding target of the inhibitors in brain tissue, we performed an *in vivo* thermal shift assay (CETSA) (*27*). The results showed a significant shift in the sEH melting curve of the hippocampus of CD-1 mice orally treated with TPPU, AS-2586114 and UB-EV-52, demonstrating *in vivo* compound-induced target stabilization, providing also evidence of central action (Fig. 2D).

### sEH inhibitors reduce biomarkers of inflammation, oxidation stress and endoplasmatic reticulum stress

Indicators of brain neuroinflammation were determined after treatment of SAMP8 mice with TPPU (5 mg/kg/day), AS-2586114 (7.5 mg/kg/day), and UB-EV-52 (5 mg/kg/day) (Fig. 3A and fig. S1). The three inhibitors reduced gene expression and protein levels of the pro-inflammatory cytokines *Il-1β, Ccl3* and, importantly, *Tnf-α* (Fig. 3B). Of note, TNF-α is not only involved in AD-related brain neuroinflammation, but also contributes to amyloidogenesis via β-secretase regulation (*28*). Additionally, these results suggest an involvement of the two main inflammasome-signaling pathways, NF-kβ and NLRP3 (*29*).

**Fig. 3.**
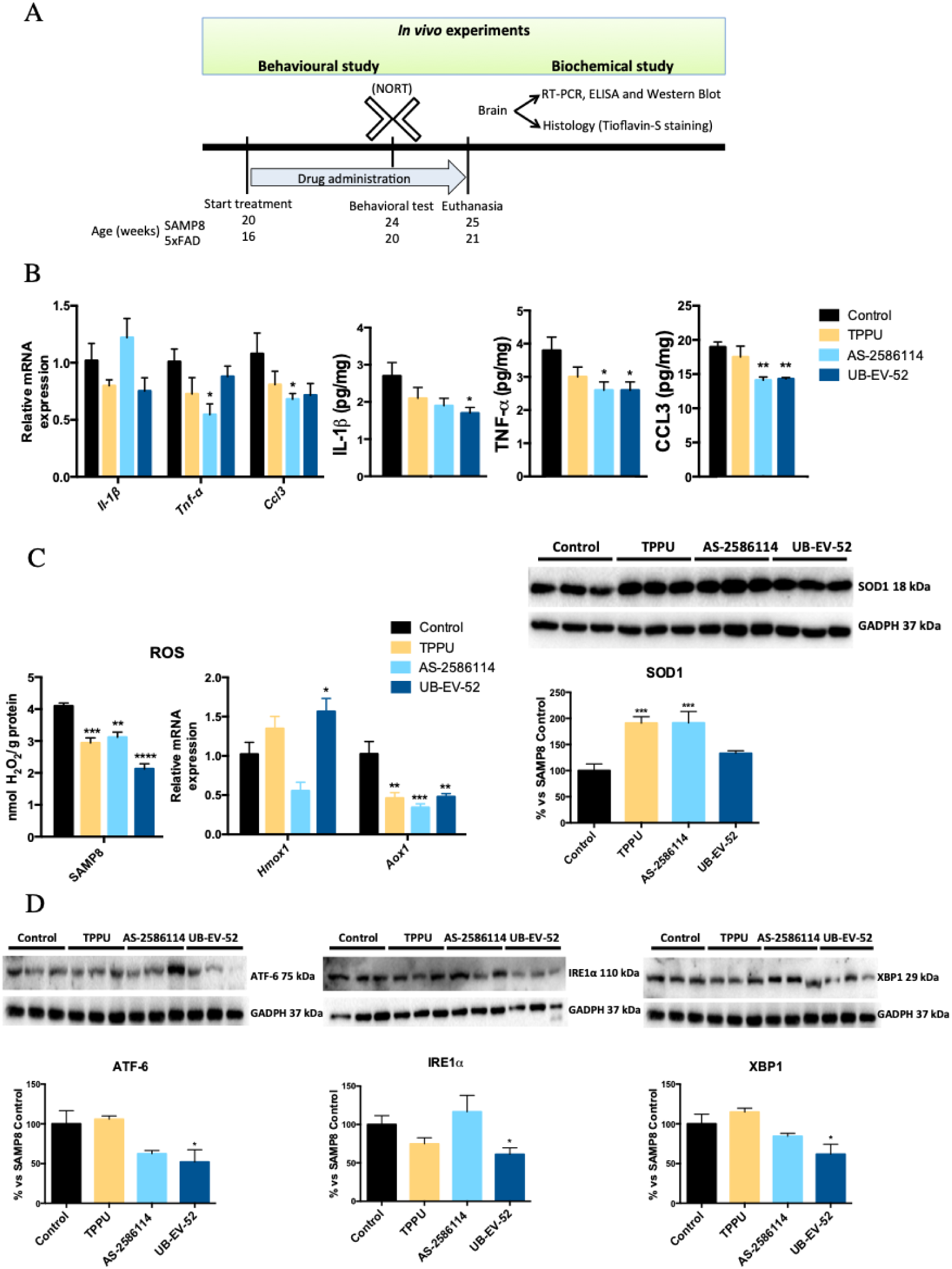
Role of sEH inhibitors in neurodegenerative biomarkers. **(A)** Scheme of experimental procedures in *in vivo* experiments. **(B)** Gene expression ofneuroinflammatory markers (*Il-1β, Tnf-α*, and *Ccl3*) and protein levels ofproinflarnmatory cytokines IL-lβ, TNF-α, and CCL3 in the hippocampus of SAMP8 mice after treatment with sEH inhibitors. **(C)** OS measured by hydrogen peroxide concentration in homogenates of the hippocampus. Representative gene expression of *Hmox*1 and *Aox*1 and representative Western blot and quantification of protein levels for (antioxidant enzyme) SOD1 in the hippocampus of SAMP8 mice after treatment with sEH inhibitors. **(D)** Representative Western blot and quantification of protein levels for ER stress markers ATF-6, IRE1α and XBP1 in the hippocampus of SAMP8 mice after treatment with sEH inhibitors. Gene expression levels were determined by real-time PCR, cytokine protein levels by ELISA and SOD1 by immunoblotting. Results are expressed as a MEAN ± SEM and were significantly different from the control group. Groups were compared by Student t-test or by One-Way ANOVA and post-hoc Dunnett’s, n = 4-6 per group, (*p<0.05, **p<0.01, ***p<0.001 and ****<0.0001) vs. Control. See partial correlations between selected variable in fig. S2 and table S4.

To investigate the influence of the sEH inhibitors in the OS process, we determined the concentration of hydrogen peroxide in the brain of SAMP8 mice. The three inhibitors significantly reduce hydrogen peroxide (Fig. 3C). Moreover, determination of the brain oxidative machinery was addressed by evaluating gene expression of *Hmox1*, *Aox1* and protein levels of SOD1 (Fig. 3C). Hmox1 (antioxidant activity) (*30*) was significantly increased by UB-EV-52 and TPPU, but not by AS-2586114 (Fig. 3C). qPCR analysis also demonstrated that treated SAMP8 mice had lower *Aox1* expression (Fig. 3C). *Aox1* controls the production of hydrogen peroxide and, under certain conditions, can catalyze the formation of superoxide (*31*). Furthermore, SOD1 (antioxidant activity) protein levels were significantly increased in all treated groups of mice (Fig. 3C), indicating a reinforcement of the antioxidant response after treatment with sEH inhibitors (*32*).

It is known that the ER stress plays a role in the pathogenesis of neurodegenerative disease, including AD (*33*), and those sEH inhibitors attenuate activation of the ER stress response (*10*). Therefore, we measured the levels of the ER stress sensors ATF-6 and IRE1α (Fig. 3D). Specially, UB-EV-52 was able to reduce the levels of both proteins. Furthermore, we evaluated XBP1, a major regulator of the unfolded protein response, which is induced by ER stress (*33*). XBP1 was significantly reduced by UB-EV-52 and slightly decreased by AS-2586114, but not by TPPU (Fig. 3D). Altogether, these results suggest that the inhibition of sEH protects against OS and the associated ER stress in the brain.

### sEH inhibitors modify the two-main physio-pathological hallmarks of AD

The brains of patients with AD contain two main physio-pathological hallmarks: tangles of hyperphosphorylated tau protein and aggregates of β-amyloid (βA). After treatment of SAMP8 and 5xFAD mice with TPPU (5 mg/kg/day), AS-2586114 (7.5 mg/kg/day), and UB-EV-52 (5 mg/kg/day), on the one hand, sEH inhibition provoked a reduction of the tau hyperphosphorylated species (Ser396 and Ser404), especially Ser404, (Fig. 4A and 4B) in agreement with the idea that OS can promote tau hyperphosphorylation and aggregation (*34,35*). On the other hand, we examined the ability of the sEH inhibitors to modify the amyloid processing cascade. While the 5xFAD transgenic mouse model develops early aggressive hallmarks of amyloid burden and cognitive loss (*36*), SAMP8 is characterized by an abnormal amyloid precursor protein (APP) processing, with a misbalance to the amyloidogenic pathway. Importantly, *C*-terminal fragments (CTF) levels are strongly implicated in neurodegeneration and the cognitive decline process in SAMP8 (*37*). We observed a substantial decrease of the ratio CTFs/APP protein levels in both mice models after treatment with sEH inhibitors (Fig. 4C and 4D). In addition, an increase of the sAPPα, and a decrease of sAPPβ supported that sEH inhibitors are able to shift the APP processing towards the non-amyloidogenic pathway, reducing then the probability of increasing βA aggregation. Finally, treatment of 5xFAD mice with sEH inhibitors had a strong effect in reducing the number of βA plaques stained with Thioflavin-S (by an average of 40%) (Fig. 4E), indicating the prevention of amyloid burden in a model characterized by βA plaque formation at early ages as two months.

**Fig. 4.**
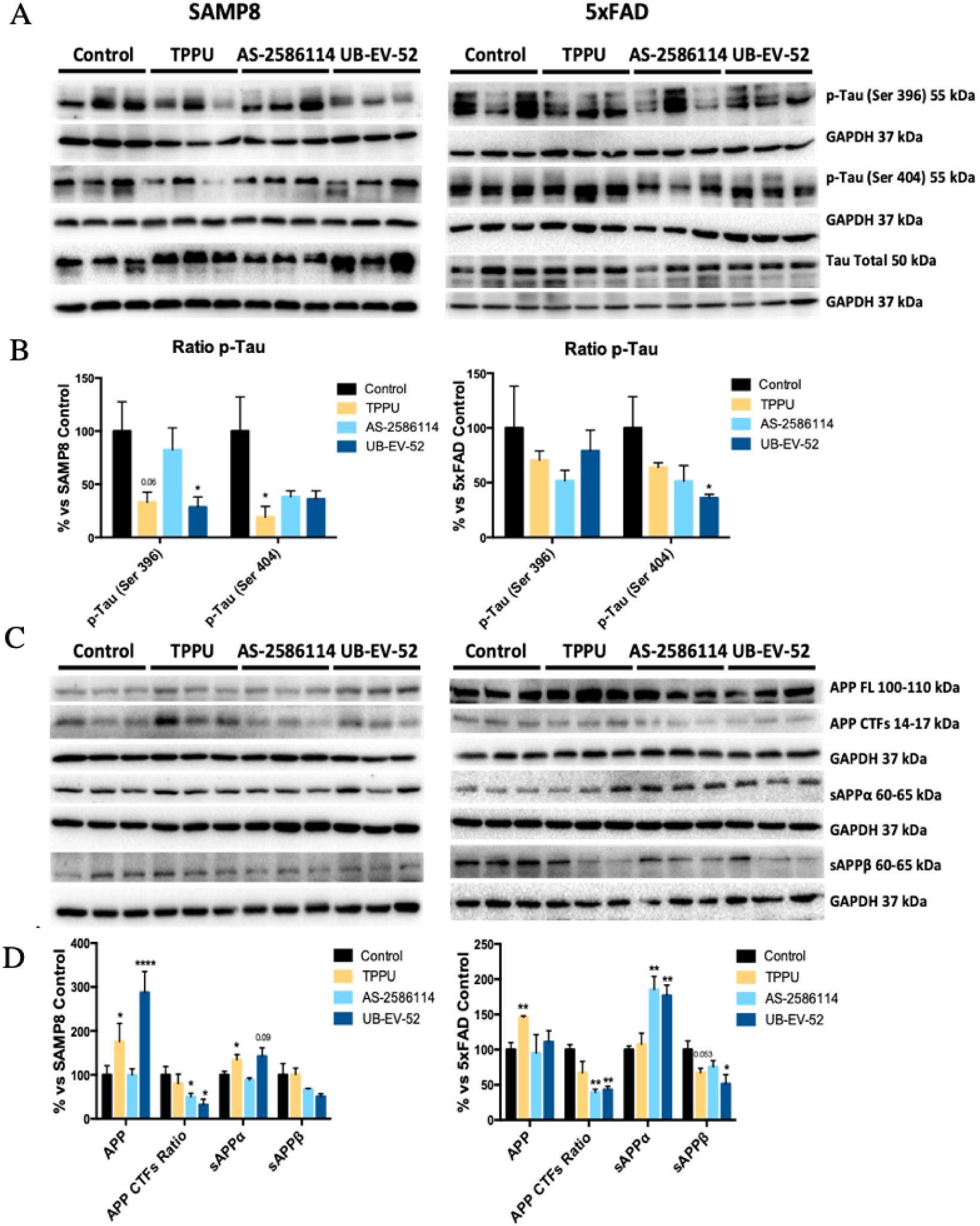
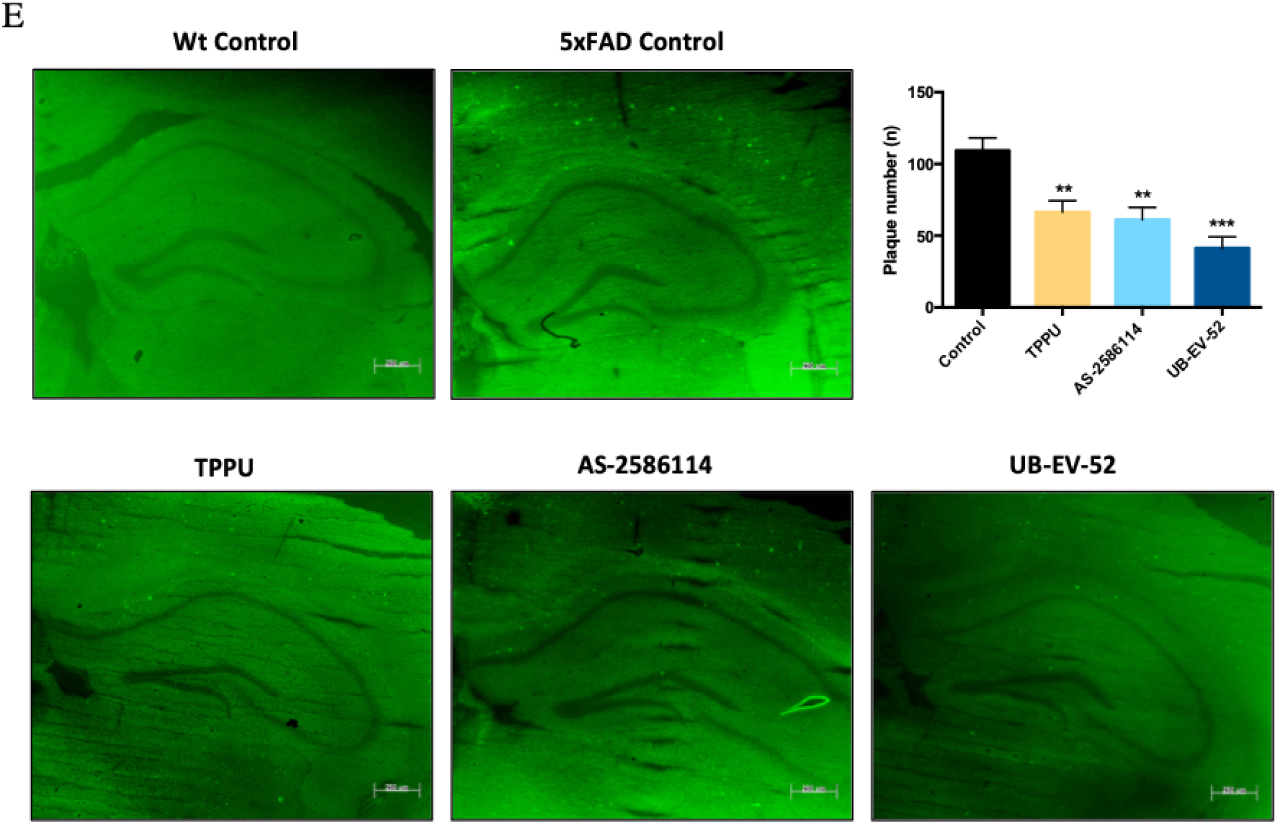
AD hallmarks in both SAMP8 and 5xFAD mice models after treatment with sEH inhibitors. **(A)** and **(B)** Representative Western blot, and quantifications for p-Tau Ser396 and p-Tau Ser404. **(C)** and **(D)** Representative Western blot, and quantifications for APP Full Length/β-CTF ratio, sAPPα and sAPPβ. **(E)** Histological images, and quantification of amyloid plaques stained with Thioflavin-S in Wt and 5xFAD. Values in bar graphs are adjusted to 100% for a protein of control group from each strain. Results are expressed as a MEAN ± SEM and were significantly different from the control group. Groups were compared by Student t-test or by One-Way ANOVA and post-hoc Dunnett’s, n = 12 per group, (*Significant at p<0.05, **Significant at p<0.01, ***Significant at p<0.001 and ****Significant at p<0.0001). See partial correlations between selected variable in fig. S2, fig. S3, table S4 and table S5.

### sEH inhibitors reduce cognitive impairment

To demonstrate the efficacy on the cognitive decline of the sEH inhibitors, a novel object recognition test (NORT) was performed to obtain a measure of cognition for short-term and long-term memory. Treatment of both murine models with the three sEH inhibitors drastically increased the Discrimination Index (DI). The significant increase indicates clear preservation of both memories (Fig. 5A and 5B). In both models, we add a comparator arm with donepezil, which is a standard of care in the treatment of AD. As expected, donepezil treatment (5 mg/kg/day) also shifted the DI to values significantly higher than zero. Remarkably, in all the conditions the sEH inhibitors reduced cognitive decline better than donepezil.

**Fig. 5.**
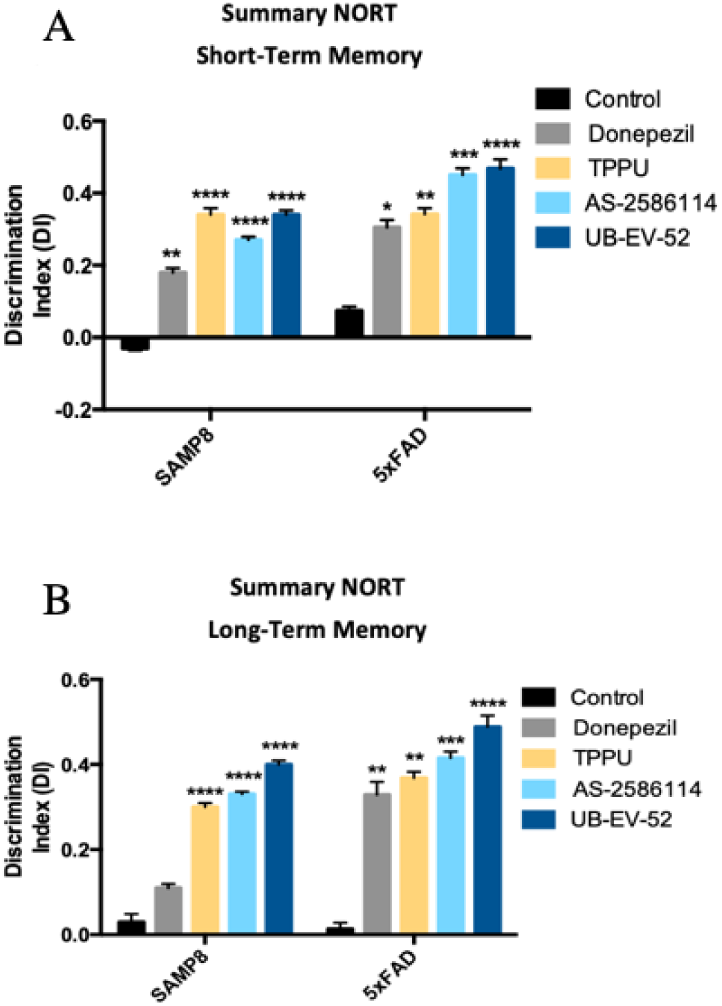
Characterization of the effect of sEH inhibitors and donepezil on cognitive status in both SAMP8 and 5xFAD mice models. **(A)** Short-term memory evaluation after 2 h acquisition trial by Discrimination Index and **(B)** Long-term memory evaluation after 24 h acquisition trial by Discrimination Index after exposure to novel objects. Results are expressed as a MEAN ± SEM and were significantly different from the control group. Groups were compared by Student t-test or by One-Way ANOVA and post-hoc Dunnett’s, n = 12 per group, (*Significant at p<0.05, **Significant at p<0.01, ***Significant at p<0.001 and ****Significant at p<0.0001). See partial correlations between selected variable in fig S2, fig. S3, table S4 and table S5.

## DISCUSSION

Our results suggest that the pharmacological stabilization of EETs in the brain has the potential to address multiples etiologies and physio-pathological processes of AD, i.e., neuroinflamation, ER stress and OS, increasing the chances of success of future therapies based on sEH inhibition. Although the positive role of sEH inhibition in multiple inflammation-related diseases has been studied, there are only a few investigations about its crucial role in the neuroinflammation process (*15, 16, 23*). In addition, the question if neuroinflammation is the malicious driver or ‘just’ a consequence still represents an important conundrum in the AD field. Our findings reinforce the idea that neuroinflammation might drive the pathogenic process in AD. A partial correlation calculation has demonstrated that the anti-inflammatory effects of sEH inhibitors correspond with changes in AD hallmarks, slowing the progression of the disease and pushing up the cognitive capabilities in the studied animal models.

While a characteristic feature of acute inflammatory processes is a general increase in the levels of classic eicosanoids (prostaglandins, leukotrienes and thromboxanes) in neurodegenerative diseases there is a basal, chronic and silent inflammation that is more related with an increase of the proinflammatory cytokines and inhibition of anti-inflammatory cytokines and by acting on different mechanisms implied in neurodegeneration (e.g., increase of the OS, increase in the glutamate pathway, etc.). This allowed us to anticipate different biological and therapeutic outcomes for sEH inhibition than for the COX and LOX pathway inhibition.

The fact that two structurally different sEH inhibitors have proven to be safe in human clinical trials for other peripheral indications (AR9281 for hypertension and GSK2256294 for diabetes mellitus, chronic pulmonary obstructive disease and subarachnoid hemorrhage) (*26, 38*) can, undoubtedly, accelerate the development of new sEH inhibitors for the treatment of AD and avoid uncertainties about the possibility of angiogenic effects when inhibiting sEH. Of relevance, for this study we have employed three structurally different sEH inhibitors, ensuring that the biological outcomes observed are not due to off-target effects related with a particular inhibitor.

In summary, we have demonstrated that sEH levels are altered in AD mice models and, more importantly, in the brain of AD patients. We have further showed that the inhibition of sEH has a plethora of central beneficial effects, such as reducing inflammation and ER stress and OS markers, p-Tau pathology and the amyloid burden and, consequently, sEH inhibitors improve the functional efficacy endpoint for cognitive status in neurodegeneration and AD animal models.

Based in the results presented in this work, we strongly believe that inhibitors of sEH could represent a completely new stand-alone treatment for the treatment of AD. However herein we do not demonstrate, and it out of the scope of this study, if inhibition of sEH could also represent an add-on therapy together with more symptomatic-like drugs i.e. donepezil.

## MATERIALS AND METHODS

### Chemicals

TPPU, 1-(1-propionylpiperidin-4-yl)-3-[4-(trifluoromethoxy)phenyl]urea, was synthesized in house as previously reported (*18*). UB-EV-52, *trans*-4-[4-(3-oxaadamant-1-yl-ureido)cyclohexyloxy]benzoic acid, was synthesized in house as previously reported (*20*). AS-2586114, 3-cyclopropyl-4-[4-[2-[[[(1*S*,2*R*,5*S*)-6,6-dimethylbicyclo[3.1.1]heptan-2-yl]methyl]amino]-2-oxoethyl]piperidin-1-yl]benzoic acid (*19*), was synthesized, as its trifluoroacetate salt, in house (see below and fig. S4).

*Synthesis of the trifluoroacetic acid salt of 3-cyclopropyl-4-*[*4-*[*2-*[[[*(1S,2R,5S)-6,6-dimethylbicyclo*[*3.1.1*]*heptan-2-yl*]*methyl*]*amino*]*-2-oxoethyl*]*piperidin-1-yl*]*benzoic acid (AS-2586114 trifluoroacetate)*.

**a)** *tert*-butyl 4-[2-[[[(*1S,2R,5S*)-6,6-dimethylbicyclo[3.1.1]heptan-2-yl]methyl]amino]-2-oxoethyl]piperidine-1-carboxylate, **1**. To a solution of (-)-*cis*-myrtanylamine (746 mg, 4.87 mmol) in EtOAc (60 mL) were added [1-(*tert*-butoxycarbonyl)piperidin-4-yl]acetic acid (1.08 g, 4.43 mmol), HOBt (897 mg, 6.64 mmol), EDC.HCl (1.27 g, 6.62 mmol) and triethylamine (1.24 mL, 8.94 mmol). The reaction mixture was stirred at room temperature for 24 h. Then, water was added (50 mL) and the phases were separated. The organic phase was washed with saturated aqueous NaHCO_3_ solution (50 mL) and brine (50 mL), dried over anh. Na_2_SO_4_, filtered and evaporated to give **1** as an orange gum (1.49 g, 89% yield). ^1^H-NMR (400 MHz, CDCl_3_) δ: 0.87 (d, *J* = 9.6 Hz, 1 H, 7’-H_c_), 1.02 [s, 3 H, C6’-(C<u>H</u>_3_)_a_], 1.10 [m, 2 H, 3(5)-H_ax_], 1.17 [s, 3 H, C6’-(C<u>H</u>_3_)_b_], 1.40-1.53 (c. s., 10 H, C(C<u>H</u>_3_)_3_ and 3’-H_a_), 1.67 [m, 2 H, 3(5)-H_eq_], 1.78-2.03 (c. s., 6 H, 4-H, 1’-H, 3’-H_b_, 4’-H_2_ and 5’-H), 2.05 (m, 2 H, NHCOC<u>H</u>_2_), 2.16 (m, 1 H, 2’-H), 2.35 (m, 1 H, 7’-H_d_), 2.70 [m, 2 H, 2(6)-H_ax_], 3.25 (m, 2 H, C<u>H</u>_2_NHCO), 4.06 [broad s, 2 H, 2(6)-H_eq_], 5.51 (m, 1 H, NH); ^13^C-NMR (100.6 MHz, CDCl_3_) δ: 19.9 (CH_2_, C3’), 23.3 [CH_3_, C6’-(<u>C</u>H_3_)_a_], 26.1 (CH_2_, C4’), 28.1 [CH_3_, C6’-(<u>C</u>H_3_)_b_], 28.5 [CH_3_, C(<u>C</u>H_3_)_3_], 32.1 [CH_2_, C3(5)], 33.3 (CH_2_, C7’), 33.6 (CH, C4), 38.8 (C, C6’), 41.5 (CH, C5’), 41.6 (CH, C2’), 43.9 [CH_2_, C2(6) and CH, C1’], 44.0 (CH_2_, NHCO<u>C</u>H_2_), 45.2 (CH_2_, <u>C</u>H_2_NHCO), 79.3 [C, O<u>C</u>(CH_3_)_3_], 155.0 (C, NCO_2_), 171.5 (C, CONH); HMRS (*m/z*): [M]^+^ calcd for [C_22_H_38_N_2_O_3_+H]^+^, 379.2955; found, 379.2967.

**b)** *N*-[[(*1S,2R,5S*)-6,6-dimethylbicyclo[3.1.1]heptan-2-yl]methyl]-2-(piperidin-4-yl)acetamide, **2**. To a solution of *tert*-butyl 4-[2-[[[(*1S,2R,5S*)-6,6-dimethylbicyclo[3.1.1]heptan-2-yl]methyl]amino]-2-oxoethyl]piperidine-1-carboxylate, **1**, (1.20 g, 3.17 mmol) in CH_2_Cl_2_ (6.5 mL), 4 N HCl in dioxane (1.85 mL) was added. The mixture was stirred at room temperature overnight. The reaction mixture was evaporated to give a residue, which was dissolved in EtOAc and washed with 2N NaOH solution. The organic phase was dried over anh. Na_2_SO_4_, filtered and evaporated to give **2** as a brown solid (0.775 g, 88% yield), mp 126-127 °C (EtOAc). ^1^H-NMR (400 MHz, CDCl_3_) δ: 0.87 (d, *J* = 9.6 Hz, 1 H, 7’-H_c_), 1.02 [s, 3 H, C6’-(C<u>H</u>_3_)_a_], 1.14 [m, 2 H, 3(5)-H_ax_], 1.17 [s, 3 H, C6’-(C<u>H</u>_3_)_b_], 1.46 (m, 1 H, 3’-H_a_), 1.69 [m, 2 H, 3(5)-H_eq_], 1.79-1.99 (c. s., 6 H, 4-H, 1’-H, 3’-H_b_, 4’-H_2_ and 5’-H), 2.04 (m, 2 H, NHCOC<u>H</u>_2_), 2.10-2.23 [c. s., 2 H, 2’-H and piperidine-N<u>H</u>), 2.34 (m, 1 H, 7’-H_d_), 2.61 [td, *J* = 12.0 Hz, *J’* = 2 Hz, 2 H, 2(6)-H_ax_], 3.04 [m, 2 H, 2(6)-H_eq_], 3.24 (m, 2 H, C<u>H</u>_2_NHCO), 5.52 (m, 1 H, NH). ^13^C-NMR (100.6 MHz, CDCl_3_) δ: 20.0 (CH_2_, C3’), 23.3 [CH_3_, C6’-(<u>C</u>H_3_)_a_], 26.1 (CH_2_, C4’), 28.1 [CH_3_, C6’-(<u>C</u>H_3_)_b_], 33.26 and 33.30 [CH_2_, C3(5) and C7’], 33.9 (CH, C4), 38.8 (C, C6’), 41.5 (CH, C5’), 41.6 (CH, C2’), 43.9 (CH, C1’), 44.7 (CH_2_, NHCO<u>C</u>H_2_), 45.2 (CH_2_, <u>C</u>H_2_NHCO), 46.6 [CH_2_, C2(6)], 171.8 (C, CONH); HMRS (*m/z*): [M]^+^ calcd for [C_17_H_30_N_2_O+H]^+^, 279.2431; found, 279.2438.

**c)** *tert*-butyl 3-bromo-4-[4-[2-[[[(*1S,2R,5S*)-6,6-dimethylbicyclo[3.1.1]heptan-2-yl]methyl]amino]-2-oxoethyl]piperidin-1-yl]benzoate, **3**. To a solution of *N*-[[(*1S,2R,5S*)-6,6-dimethylbicyclo[3.1.1]heptan-2-yl]methyl]-2-(piperidin-4-yl)acetamide, **2**, (734 mg, 2.64 mmol) in anh. DMSO (9 mL) a solution of *tert*-butyl 3-bromo-4-fluorobenzoate (727 mg, 2.64 mmol) in anh. DMSO (4 mL) was added. Then, anh. K_2_CO_3_ (729 mg, 5.28 mmol) was added and the mixture was heated at 90 °C for 36 h. The brown suspension was poured into cold water (50 mL) and extracted with EtOAc (3 x 50 mL). The organic phase was washed with water (50 mL), brine (50 mL), dried over anh. Na_2_SO_4_, filtered and evaporated to give a residue which was purified by column chromatography (hexane/ethyl acetate mixtures) to give **3** as a white solid (0.57 g, 40% yield), mp 85-86 °C (CH_2_Cl_2_); ^1^H-NMR (400 MHz, CDCl_3_) δ: 0.89 (d, *J* = 9.6 Hz, 1 H, 7’-H_c_) 1.04 [s, 3 H, C6’-(C<u>H</u>_3_)_a_], 1.19 [s, 3 H, C6’-(C<u>H</u>_3_)_b_], 1.42-1.55 [c. s., 3 H, 3(5)-H_ax_ and 3’-H_a_], 1.57 [s, 9 H, C(C<u>H</u>_3_)_3_], 1.83 [m, 2 H, 3(5)-H_eq_], 1.87-2.08 [c. s., 6 H, 4-H, 1’-H, 3’-H_b_, 4’-H_2_ and 5’-H], 2.14 (m, 2 H, NHCOC<u>H</u>_2_), 2.18 (m, 1 H, 2’-H), 2.36 (m, 1 H, 7’-H_d_), 2.69 [m, 2 H, 2(6)-H_ax_], 3.28 (m, 2 H, C<u>H</u>_2_NHCO), 3.43 [m, 2 H, 2(6)-H_eq_], 5.51 (m, 1 H, NH), 6.99 (d, *J* = 8.4 Hz, 1 H, 5’’-H), 7.85 (dd, *J* = 8.4 Hz, *J’* = 2 Hz, 1 H, 6’’-H), 8.13 (d, *J* = 2.0 Hz, 1 H, 2’’-H); ^13^C-NMR (100.6 MHz, CDCl_3_) δ: 20.0 (CH_2_, C3’), 23.3 [CH_3_, C6’-(<u>C</u>H_3_)_a_], 26.1 (CH_2_, C4’), 28.1 [CH_3_, C6’-(<u>C</u>H_3_)_b_], 28.3 [CH_3_, C(<u>C</u>H_3_)_3_], 32.4 [CH_2_, C3(5)], 33.2 (CH, C4), 33.3 (CH_2_, C7’), 38.8 (C, C6’), 41.5 (CH, C5’), 41.6 (CH, C2’), 44.0 (CH, C1’), 44.1 (CH_2_, NHCO<u>C</u>H_2_), 45.3 (CH_2_, <u>C</u>H_2_NHCO), 52.1 [CH_2_, C2(6)], 81.2 [C, O<u>C</u>(CH_3_)_3_], 118.7 (C, C3’’), 120.1 (CH, C5’’), 127.2 (C, C1’’), 129.7 (CH, C6’’), 135.2 (CH, C2’’), 155.1 (C, C4’’), 164.7 (C, NCO_2_), 172.0 (C, CONH); HMRS (*m/z*): [M]^+^ calcd for [C_29_H_41_BrN_2_O_3_+H]^+^, 533.2373; found, 533.2372.

**d)** *tert*-butyl 3-cyclopropyl-4-[4-[2-[[[(*1S,2R,5S*)-6,6-dimethylbicyclo[3.1.1]heptan-2-yl]methyl]amino]-2-oxoethyl]piperidin-1-yl]benzoate, **4**. A mixture of *tert*-butyl 3-bromo-4-[4-[2-[[[(*1S,2R,5S*)-6,6-dimethylbicyclo[3.1.1]heptan-2-yl]methyl]amino]-2-oxoethyl]piperidin-1-yl]benzoate, **3**, (546 mg, 1.02 mmol), cyclopropylboronic acid (176 mg, 2.05 mmol), tetrakis(triphenylphosphine) palladium(0) (118 mg, 0.102 mmol) and K_3_PO_4_ (649 mg, 3.06 mmol) in 1,4-dioxane (11 mL) was heated at 100 °C for 9 hours. The mixture was filtered with Celite^®^ and eluted by EtOAc. Evaporation under vacuum of the organic phase gave a residue which was purified by column chromatography (hexane/ethyl acetate mixtures) to give **4** as a foaming white solid (381 mg, 75% yield), mp 88-89 °C (CH_2_Cl_2_); ^1^H-NMR (400 MHz, CDCl_3_) δ: 0.78 (m, 2 H, cyclopropyl-C<u>H</u>_2_), 0.89 (d, *J* = 9.6 Hz, 1 H, 7’-H_c_), 0.99 (m, 2 H, cyclopropyl-C<u>H</u>_2_), 1.04 [s, 3 H, C6’-(C<u>H</u>_3_)_a_], 1.19 [s, 3 H, C6’-(C<u>H</u>_3_)_b_], 1.37 [c. s., 3 H, 3(5)-H_ax_ and 3’-H_a_], 1.56 [s, 9 H C(C<u>H</u>_3_)_3_], 1.84 [m, 2 H, 3(5)-H_eq_], 1.87-2.07 [c. s., 6 H, 4-H, 1’-H, 3’-H_b_, 4’-H_2_ and 5’-H], 2.10-2.26 (c. s., 4 H, NHCOC<u>H</u>_2_, 2’-H and cyclopropyl-C<u>H</u>), 2.36 (m, 1 H, 7’-H_d_), 2.70 [td, *J* = 11.8 Hz, *J’* = 1.8 Hz, 2 H, 2(6)-H_ax_], 3.28 (m, 2 H, C<u>H</u>_2_NHCO), 3.41 [m, 2 H, 2(6)-H_eq_], 5.50 (m, 1 H, NH), 6.94 (d, *J* = 8.4 Hz, 1 H, 5’’-H), 7.40 (d, *J* = 2.0 Hz, 1 H, 2’’-H). 7.71 (dd, *J* = 8.4 Hz, *J’* = 2 Hz, 1 H, 6’’-H); ^13^C-NMR (100.6 MHz, CDCl_3_) δ: 9.6 (<u>C</u>H_2_, cyclopropyl), 11.4 (<u>C</u>H, cyclopropyl), 20.0 (CH_2_, C3’), 23.3 [CH_3_, C6’-(<u>C</u>H_3_)_a_], 26.1 (CH_2_, C4’), 28.1 (CH_3_, C6’-(<u>C</u>H_3_)_b_], 28.4 [CH_3_, C(<u>C</u>H_3_)_3_], 32.7 (CH_2_) and 32.8 (C3 and C5), 33.3 (CH_2_, C7’), 33.5 (CH, C4), 38.8 (C, C6’), 41.5 (CH, C5’), 41.6 (CH, C2’), 44.0 (CH, C1’), 44.2 (CH_2_, NHCO<u>C</u>H_2_), 45.3 (CH_2_, <u>C</u>H_2_NHCO), 52.3 [CH_2_, C2(6)], 80.5 [C, O<u>C</u>(CH_3_)_3_], 117.9 (CH, C5’’), 125.2 (CH, C2’’), 125.8 (C, C1’’), 127.5 (CH, C6’’), 136.7 (C, C3’’), 156.6 (C, C4’’), 166.2 (C, NCO_2_), 171.8 (C, CONH); HMRS (*m/z*): [M]^+^ calcd for [C_31_H_46_N_2_O_3_+H]^+^, 495.3581; found, 495.3584.

**e)** AS-2586114 (as trifluoroacetate salt). To a solution of *tert*-butyl 3-cyclopropyl-4-[4-[2-[[[(*1S,2R,5S*)-6,6-dimethylbicyclo[3.1.1]heptan-2-yl]methyl]amino]-2-oxoethyl]piperidin-1-yl]benzoate, **4**, (338 mg, 0.68 mmol) in CH_2_Cl_2_ (4.2 mL), TFA was added (2.52 mL) and the reaction mixture was stirred at room temperature overnight. The mixture was evaporated to give an orange gum. After the addition of diethyl ether, the salt precipitated as a pale white yellowish solid (254 mg, 68% yield), mp 139-140 °C (Et_2_O); [α]^D^ 20 (deg cm^3^ g^-1^ dm^-1^) = −7.0 (c = 1.0 g cm^-3^ in methanol); ^1^H-NMR (400 MHz, DMSO) δ: 0.69 (m, 2 H, cyclopropyl-C<u>H</u>_2_), 0.83 (d, *J* = 9.6 Hz, 1 H, 7’-H_c_), 0.96-1.06 [c. s., 5 H, cyclopropyl-C<u>H</u>_2_ and C6’-(C<u>H</u>_3_)_a_], 1.15 [s, 3 H, C6’-(C<u>H</u>_3_)_b_], 1.28-1.48 [c. s., 3 H, 3(5)-H_ax_ and 3’-H_a_], 1.74 [m, 2 H, 3(5)-H_eq_], 1.77-1.94 (c. s., 6 H, 4-H, 1’-H, 3’-H_b_, 4’-H_2_ and 5’-H), 2.0-2.18 (c. s., 4 H, NHCOC<u>H</u>_2_, 2’-H and cyclopropyl-C<u>H</u>), 2.31 (m, 1 H, 7’-H_d_), 2.67 [m, 2 H, 2(6)-H_ax_], 3.06 (m, 2 H, C<u>H</u>_2_NHCO), 3.34 (m, 2 H, 2(6)-H_eq_), 4.90 (broad s, NH^+^), 7.04 (d, *J* = 8.4 Hz, 1 H, 5’’-H), 7.30 (d, *J* = 2.0 Hz, 1 H, 2’’-H), 7.67 (dd, *J* = 8.4 Hz, *J’* = 2.0 Hz, 6’’-H), 7.80 (m, 1 H, NH), the signal of the Ar-COO<u>H</u> is not observed; ^13^C-NMR (100.6 MHz, DMSO) δ: 9.46 (<u>C</u>H_2_, cyclopropyl), 11.1 (<u>C</u>H, cyclopropyl), 19.2 (CH_2_, C3’), 22.9 [CH_3_, C6’-(<u>C</u>H_3_)_a_], 25.7 (CH_2_, C4’), 27.8 [CH_3_, C6’-(<u>C</u>H_3_)_b_], 32.0 [CH_2_, C3(5)], 32.7 (CH_2_, C7’), 32.8 (CH, C4), 38.2 (C, C6’), 40.7 (CH, C5’), 40.8 (CH, C2’), 42.5 (CH_2_, NHCO<u>C</u>H_2_), 43.0 (CH, C1’), 44.0 (CH_2_, <u>C</u>H_2_NHCO), 51.8 [CH_2_, C2(6)], 115.2 (C, *J* = 289.7 Hz, CF_3_), 118.1 (CH, C5’’), 124.3 (C, C1’’), 124.6 (CH, C2’’), 127.4 (CH, C6’’), 136.1 (C, C3’’), 156.2 (C, C4’’), 158.3 (C, *J* = 38.2 Hz, CF_3_-<u>C</u>OO^-^), 167.3 (C, COOH), 170.8 (C, CONH); IR (ATR) ν: 835, 939, 1019, 1071, 1232, 1299, 1384, 1403, 1431, 1464, 1497, 1559, 1601, 1650, 1682, 1696, 2742, 2813, 2865, 2922, 2984, 3078, 3293 cm^-1^; elemental analysis: calcd for C_29_H_39_F_3_N_2_O_5_: C 63.03%, H 7.11%, N 5.07%; found: C 63.33%, H 7.30%, N 4.92%.

### Animals

The Senescence-Accelerated Prone Mouse 8 (SAMP8), a non-transgenic mouse, was established through phenotypic selection of the AKR/J mice strain and is an attractive model to study aging processes and, specially, age-related deterioration in learning and memory, emotional disorders and neurochemical alterations around 5 months of age (*39-41*). Some of the changes in the mice are directly related to AD such as APP processing alteration and Tau hyperphosphorylation (*42*). Moreover, these animals manifest evidence of early alterations in OS and inflammation (*43*).

The 5xFAD is a double transgenic APP/PS1 that co-expresses five mutations of AD and that rapidly develops severe amyloid pathology with high levels of intraneuronal Aβ42 around 2 months of age. The model was generated by the introduction of human APP with the Swedish mutations (K670N/M671L), Florida (I716V), London (Val717Ile) and the introduction of PS1 M146L and L286V. Moreover, 5xFAD presents neuronal loss and cognitive deficits in spatial learning (at approximately four to five months) (*44*). Tangles are not typical in this model. Long-term potentiation (LTP) is normal in young animals, but becomes impaired around six months (*45*).

20-Week old male SAMP8 mice (n = 48) and 16-week-old male 5xFAD mice (n = 48) were used to carry out cognitive and molecular analyses. The animals were randomly allocated to experimental groups. We divided these animals into eight groups: SAMP8 Control (n = 12) and 5xFAD Control (n = 12), animals administered with vehicle (2-hydroxypropyl)-β-cyclodextrin 1.8%, and both strains treated with different sEH inhibitors: TPPU (TPPU, n = 12) at 5 mg/Kg/day, AS-2586114 (AS-2586114, n = 12) at 7.5 mg/Kg/day and UB-EV-52 (UB-EV-52, n = 12) at 5 mg/Kg/day. sEH inhibitors were administered through drinking water for 4 weeks diluted in 1.8% (2-hydroxypropyl)-β-cyclodextrin. Water consumption was controlled each week and the sEH inhibitor concentration was adjusted accordingly to reach the precise dose.

Animals had free access to food and water, and were kept under standard temperature conditions (22 ± 2 °C) and 12 hours: 12 hours light-dark cycles (300 lux/0 lux).

Studies and procedures involving mouse brain dissection and extractions were performed in accordance with the institutional guidelines for the care and use of laboratory animals and approved by the Ethics Committee for Animal Experimentation at the University of Barcelona.

### Novel Object Recognition Test

This test allows evaluating short- and long-term recognition memory involving cortical areas and the hippocampus (*46*). The experimental apparatus used for this test was a 90-degree, two-arm, 25-cm-long, 20-cm-high maze of black polyvinyl chloride. Light intensity in the middle of the field was 30 lux. First, mice were individually habituated to the apparatus for 10 min per day during 3 days. On the day four, the animals were allowed to freely explore 10 min acquisition trial (First trial), during which they were placed in the maze in the presence of two identical novel objects (A and A or B and B) placed at the end of each arm. The mouse was then removed from the apparatus and returned to its home cage. A 10-min retention trial (second trial) was carried out 2 hours (short-term memory) or 24 hours (long-term memory) later. During this second trial, objects A and B were placed in the maze, and the times that the animal took to explore the novel object (TN) and the old object (TO) were recorded. 24 hours after the acquisition trial, the mice were tested again, with a new object and an object identical to the new one in the previous trial (A and C or B and C). The time that mice explored the TN and time that mice explored the TO were measured from the video recordings from each trial session. A Discrimination Index (DI) was defined as (TN-TO)/(TN+TO). Exploration of an object by a mouse was defined as pointing the nose towards the object at a distance ≤ 2 cm and/or touching it with the nose. Turning or sitting around the object was not considered exploration. In order to avoid object preference biases, objects A and B were counterbalanced so that one half of the animals in each experimental group were first exposed to object A and then to object B, whereas the other half first saw object B and then object A. The maze, the surface, and the objects were cleaned with 70% ethanol between the animals’ trials so as to eliminate olfactory cues.

### Biochemical experiments

#### Murine brain tissue preparation

SAMP8 mice were euthanized 3 days after the behavioural test completion by cervical dislocation. Brains were immediately removed from the skull. Cortex and hippocampus were then isolated and frozen on powdered dry ice. They were maintained at −80 °C for biochemical experiments.

5xFAD mice were anesthetized with pentobarbital 3 days after behavioral test completion and transcardially perfused with saline. Afterward, brains were dissected. One hemisphere was post-fixed overnight in 4% PFA with 15% sucrose at 4 °C, while the other hemisphere was dissected into cortex and hippocampus, frozen on powdered dry ice and maintained at −80 °C for further biochemical experiments.

### Human cases

Tissue samples were obtained from the Institute of Neuropathology-IDIBELL Brain Bank, Hospitalet de Llobregat following the guidelines of Spanish legislation on this matter and the approval of the local ethics committee. The interval between death and autopsy was between 1 to 10 hours (fig. S1). The brain tissue was immediately frozen on metal plates over dry ice, placed in individual air-tight plastic bags, and maintained at −80 °C for biochemical experiments. Neuropathologic diagnosis of AD was based on the classification of Braak and Braak (*47,48*).

### Protein levels determination by Western blotting

For Western blotting (WB), aliquots of 15 µg of hippocampal protein were extracted with lysis buffer containing phosphatase and protease inhibitors (Cocktail II, Sigma) and protein concentration was determined by Bradford’s method. Protein samples from 24 mice of both strains (n = 3 per group) were separated by SDS-PAGE (8-20%) and transferred onto PVDF membranes (Millipore). Afterward, membranes were blocked in 5% non-fat milk in 0.1% Tween20-TBS (TBS-T) for 1 hour at room temperature, followed by overnight incubation at 4 °C with the primary antibodies, that are presented in Table 1.

**Table 1.**
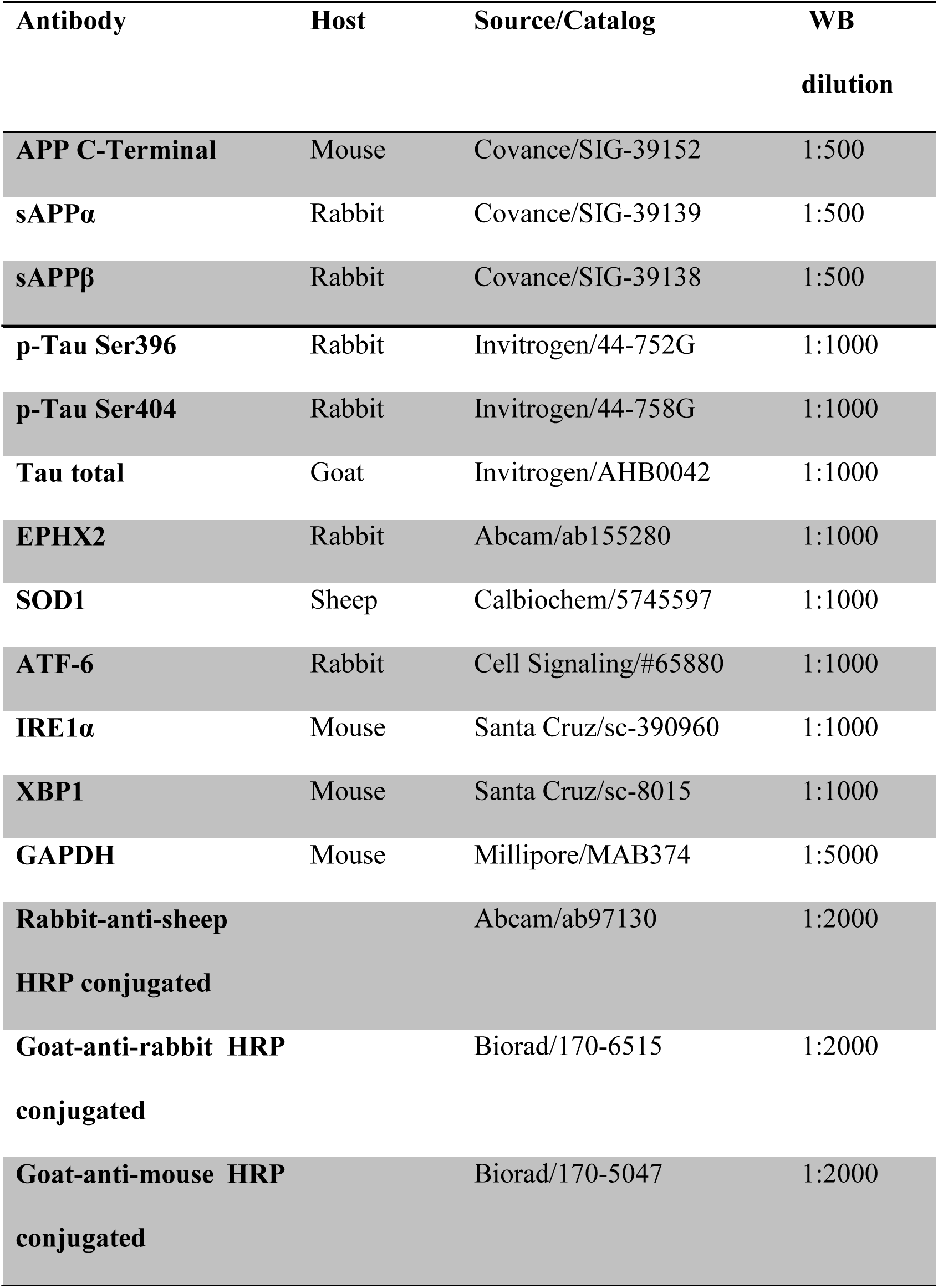
Antibodies used in Western blotting.

The next day, membranes were washed with TBS-T and incubated with secondary antibodies for 1 hour at room temperature. Immunoreactive proteins were viewed with a chemiluminescence-based detection kit, following the manufacturer’s protocol (ECL Kit, Millipore) and digital images were acquired using a ChemiDoc XRS+System (BioRad). Semi-quantitative analyses were carried out using ImageLab Software (BioRad) and results were expressed in Arbitrary Units (AU), considering the control mice group as 100%. Protein loading was routinely monitored by immunodetection of GAPDH.

### Protein levels determination by ELISA

Immunoreactive IL-1β, TNF-α, and CCL3 protein levels from 40 mice of both strains (n = 4-5 per group) were determined using mouse-specific ELISA kits according to manufacturer’s instructions (Mouse IL-1β ELISA Ready-Set-Go, eBioscience, 88-7013-77; Mouse TNF-α Ready-Set-Go, eBioscience, 88-7324-77; Mouse MIP-1 alpha (CCL3) Ready-SET-Go, eBioscience, 88-56013-22) on brain tissue samples from control mice and treated mice with different sEH inhibitors. Brain samples were placed in sterile phosphate-buffered saline (PBS) containing a protease inhibitor cocktail, homogenized, centrifuged, and, finally, the supernatant was removed. The latter was diluted in sterile PBS buffer and assayed for IL-1β and TNF-α by mouse specific sandwich ELISA. Equivalent amounts of proteins were used for the analyses. Cytokine levels were expressed as pg/mg of protein.

### RNA extraction and gene expression determination

Total RNA isolation was carried out using TRIzol® reagent according to manufacturer’s instructions. The yield, purity, and quality of RNA were determined spectrophotometrically with a NanoDrop^™^ ND-1000 (Thermo Scientific) apparatus and an Agilent 2100B Bioanalyzer (Agilent Technologies). RNAs with 260/280 ratios and RIN higher than 1.9 and 7.5, respectively, were selected. Reverse Transcription-Polymerase Chain Reaction (RT-PCR) was performed as follows: 2 μg of messenger RNA (mRNA) was reverse-transcribed using the High Capacity cDNA Reverse Transcription Kit (Applied Biosystems). Real-time quantitative PCR (qPCR) from 48 mice of both strains (n = 4-6 per group) was used to quantify mRNA expression of OS and inflammatory genes.

SYBR® Green real-time PCR was performed in a Step One Plus Detection System (Applied-Biosystems) employing SYBR® Green PCR Master Mix (Applied-Biosystems). Each reaction mixture contained 7.5 μL of complementary DNA (cDNA) (which concentration was 2 μg), 0.75 μL of each primer (which concentration was 100 nM), and 7.5 μL of SYBR® Green PCR Master Mix (2X).

TaqMan-based real-time PCR (Applied Biosystems) was also performed in a Step One Plus Detection System (Applied-Biosystems). Each 20 μL of TaqMan reaction contained 9 μL of cDNA (25 ng), 1 μL 20X probe of TaqMan Gene Expression Assays and 10 μL of 2X TaqMan Universal PCR Master Mix.

Data were analyzed utilizing the comparative Cycle threshold (Ct) method (ΔΔCt), where the housekeeping gene level was used to normalize differences in sample loading and preparation (*49*). Normalization of expression levels was performed with β*-actin* for SYBR^®^ Green-based real-time PCR and TATA-binding protein *(Tbp)* for TaqMan-based real-time PCR. Primers sequences and TaqMan probes used in this study are presented in Table 2. Each sample was analyzed in duplicate, and the results represent the n-fold difference of the transcript levels among different groups.

**Table 2.**
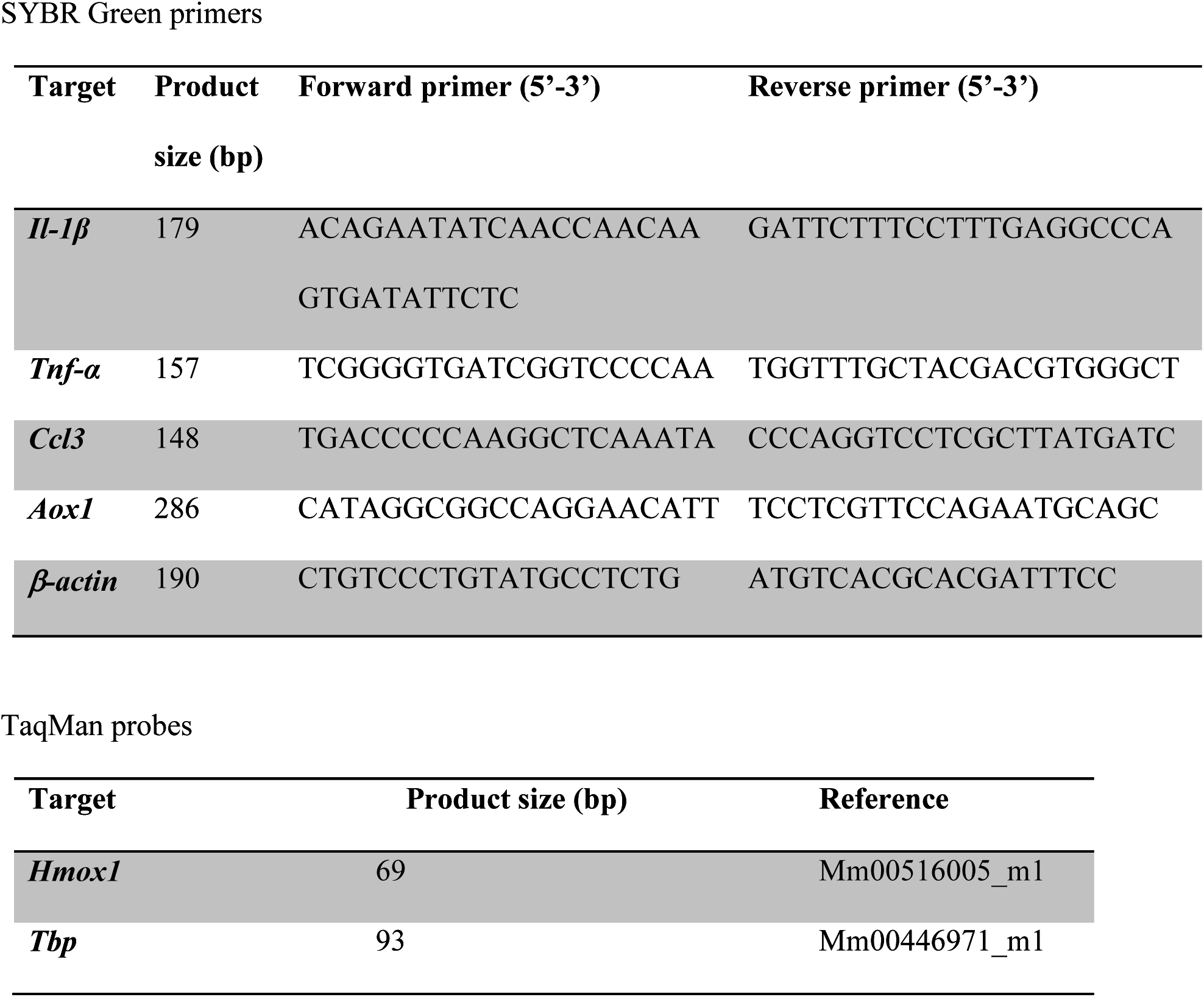
Primers and probes used in qPCR studies.

### Aβ plaque histology

Amyloid plaques from 16 5xFAD mice (n = 4 per group) were stained with Thioflavin-S. The frozen brains were embedded into OCT Cryostat Embedding Compound (Tissue-Tek, Torrance, CA, USA) and then cut into 20 µm- thick sections at −20 °C using a cryostat (Leica Microsystems, Germany) and placed on slides. For the Thioflavin-S staining procedure, the brain sections were first rehydrated at room temperature by 5 min incubation in PBS. To continue with, brain sections were incubated with 0.3% Thioflavin-S (Sigma-Aldrich) solution for 20 min at room temperature in the dark. Subsequently, these samples were submitted to washes in 3 min series, specifically with two washes using 80% ethanol, one wash using 90% ethanol and three washes with PBS. Then, slides were mounted with Fluoromount-G^TM^ (EMS, Hatfield, NJ, USA) and allowed to dry overnight. Image acquisition was performed with a fluorescence laser microscope (Olympus BX51; Germany). For plaque quantification, similar and comparable histological areas were selected, focusing on the adjacent positioning of the whole cortical area and the hippocampus.

### Determination of oxidative stress

Hydrogen peroxide from 40 hippocampus samples of mice of both strains (n = 5 per group) was measured as an indicator of oxidative stress and it was quantified using the Hydrogen Peroxide Assay Kit (Sigma-Aldrich, St. Louis, MI) according to the manufacturer’s instructions.

### Thermal Shift Assay (CETSA) for *in vivo* brain tissue

The Thermal Shift Assay (TSA) was carried out based on a previously described protocol with modifications (*27,50,51*). For the *in vivo* hippocampus tissue TSA experiment, male CD-1 mice 8-week-old (35-45 g; n = 3 per group) were treated with a single dose of each sEH inhibitor at 5 mg/Kg for TPPU, 7.5 mg/Kg for AS-2586114 and 5 mg/Kg for UB-EV-52, by intraperitoneal injection (IP) in 20% of (2-hydroxypropyl)-β-cyclodextrin, and, after one hour and a half, mice were euthanized by cervical dislocation. Immediately, brains were removed from the skull, frozen on the powdered dry ice and were maintained at −80 °C.

The brains from the different above-mentioned groups were homogenized in 200 μL PBS with EDTA 5 mM and complete protease inhibitor cocktail II (Sigma). The homogenates were aliquoted into microtubes for each treatment group and subjected to 7-point temperature gradient per duplicate (room temperature, 44 °C, 48 °C, 52 °C, 56 °C, 60 °C, and 64 °C) for 8 min using a VeriFlex temperature control technology of the thermocycler (Applied Biosystems Veriti^TM^ Thermal Cycler). Immediately, homogenates were incubated at room temperature (25 °C water bath) for 5 min and lysed with freeze-thawed three times using liquid nitrogen. Then, the supernatant fraction for each 7-point temperature homogenate from each treatment group was separated from the cellular debris and protein aggregates by centrifugation at 20,000 g for 20 min at 4 °C, and the duplicate of 7-point temperature gradient without centrifugation were the whole fraction. All these 14 samples for each treatment were transferred to new microtubes and analysed by sodium dodecyl sulfate polyacrylamide gel electrophoresis (SDS-PAGE) followed by western blotting analysis by using rabbit monoclonal anti-EPHX2 antibody as described above. The appropriate temperatures were determined in preliminary TSA experiments (Data not shown).

### Data analysis

The statistical analysis was conducted using GraphPad Prism ver.6 statistical software. Data were expressed as the mean ± Standard Error of the Mean (SEM) from at least 3 samples per group. Means were compared to One-Way ANOVA analysis of variance, followed by Dunnett’s post-hoc analysis. Comparison between groups were also performed by two-tail Student’s *t*-test for independent samples. Statistical significance was considered when p-values were <0.05. The statistical outliers were carried out with Grubss’ test and subsequently removed from the analysis. Behavioral analyses were performed blindly.

In addition, partial correlation controlling for each strain was calculated using SPSS 21.00, between the following variables: DI 2 h, DI 24 h, p-Tau Ser396, p-Tau Ser404, reactive oxygen species (ROS), *Aox1*, *Hmox*1, SOD1, *Il-1β*, *Tnf-α*, *Ccl3*, IL-1β, TNF-α, CCL3, ATF-6, IRE1α, XBP1, APP, sAPPα, sAPPβ and β-CTF for SAMP8 mice model. DI 2 h, DI 24 h, p-Tau Ser396, p-Tau Ser404, APP, sAPPα, sAPPβ, β-CTF and amyloid plaques for 5xFAD mice model. Partial correlation coefficients were calculated using the data from behavioural parameters from NORT; protein levels for SOD1, p-Tau Ser396 and 404, IL-1β, TNF-α, CCL3, APP, sAPPα, sAPPβ and β-CTF; ROS for hydrogen peroxide concentration; gene expression for *Aox1*, *Hmox1, Il-1β*, *Tnf-α* and *Ccl3*; and number of amyloid plaques obtained with Thioflavin-S staining.

### Microsomal stability of human, rat and mice microsomes

The human, rat and mice microsomes employed were purchased from Tebu-Xenotech. The compound was incubated at 37 °C with the microsomes in a 50 mM phosphate buffer (pH = 7.4) containing 3 mM MgCl_2_, 1 mM NADP, 10 mM glucose-6-phosphate and 1 U/mL glucose-6-phosphate-dehydrogenase. Samples (75 µL) were taken from each well at 0, 10, 20, 40 and 60 min and transferred to a plate containing 4 °C 75 µL acetonitrile and 30 µL of 0.5% formic acid in water were added for improving the chromatographic conditions. The plate was centrifuged (46000 g, 30 min) and supernatants were taken and analyzed in a UPLC-MS/MS (Xevo-TQD, Waters) by employing a BEH C18 column and an isocratic gradient of 0.1% formic acid in water: 0.1% formic acid acetonitrile (60:40). The metabolic stability of the compounds was calculated from the logarithm of the remaining compounds at each of the time points studied.

### Cytochrome inhibition

The human cytochrome isoforms were purchased from Corning unless CYP2C19 that was purchased from Cypex. Compounds were incubated for 5 min at 37 °C in 50 mM phosphate buffer (pH = 7.4) in the presence of 1.3 mM NADP (0.0082 mM for CYP2D6), 3.3 mM MgCl_2_ (0.41 mM for CYP2D6) and 0.4 U/mL Glucose-6-phosphate Dehydrogenase. After this time an enzyme/substrate mix was added to each well. This mix contained the different human cytochrome P450 isoforms and the specific substrates for each one: 3-cyano-7-ethoxycoumarin for CYP1A2 (5 µM) and CYP2C19 (25 µM), 75 µM 7-methoxy-4-trifluoromethylcoumarin for CYP2C9, 1.5 µM 3-[2-(N,N-diethyl-N-methylammonium)-ethyl]-7-methoxy-4-methylcoumarin iodide for CYP2D6, and 50 µM 7-benzyloxytrifluoromethyl coumarin (BFC) or 1 µM dibenzylfluorescein (DBF) for CYP3A4. Furafyline, sulfaphenazole, tranylcypromine and ketoconazole were employed as reference compounds in the assays. The incubation time varied between the different isoforms (15 min for CYP1A2, 45 min for CYP2C9 and CYP2C19, 30 min for CYP3A4 with BFC and CYP2D6 and 10 min for CYP3A4 with DBF) and the reaction was stopped by adding a stop solution containing 80% acetonitrile, 20% 0.5 M Tris-HCl for all the isoforms unless for CYP3A4 with DBF where a 2 N NaOH solution was employed as stop solution. Fluorescence due to the substrate metabolism was detected in a M1000Pro reader (Tecan) and the percentage of inhibition was calculated from the activity shown in the absence of inhibitor.

### hERG ion channel inhibition

hERG channel inhibitory activity was performed by automated patch-clamp on an automated patch clamp (Ionflux, Fluxion). CHO-hERG cell line was purchased from B’Sys. Cells were grown in F-12 Ham supplemented with Glutamax, 10% FBS, 1% penicillin/streptomycin, 100 µg/mL hygromycin and 100 µg/mL neomycin at 37 °C in an atmosphere of 5% CO_2_. 48 h prior to assay the cells were incubated at 30 °C in a 5% CO_2_ atmosphere in order to increase channel expression. On the day of the assay the cells were suspended in extracellular solution (2 mM CaCl_2_, 1 mM MgCl_2_, 10 mM HEPES, 4 mM KCl, 145 mM NaCl, 10 mM glucose, pH = 7.4, 305-320 mOsm) at a concentration of 3·10^6^ cells/mL. The compound was added to a 384-well Ionflux population patch-clamp plate and the intracellular solution (5.374 mM CaCl_2_, 1.75 mM MgCl_2_, 10 mM HEPES, 10mM EGTA, 120 mM KCl, 4 mM sodium ATP, pH = 7.2, 295-310 mOsm) in the wells. The cells (80 μL/well) were added to the plate and the hERG activity was registered beginning with a 5 seconds depolarization phase (from −80 mV to +20 mV) followed by a 5-second tail at - 50 mV and a repolarization at −80 mV. This cycle was repeated every 15 seconds. Current inhibition was calculated from Ionflux Data Analyzer software (V4.6) from the basal activity of the channel.

### Solubility

A 10 mM stock solution of the compound was serially diluted in 100% DMSO and 2.5 µL of this solution was added to a 384-well UV-transparent plate (Greiner) containing 47.5 µL of PBS. The plate was incubated at 37 °C for 4 h and the light scattering was measured in a Nephelostar Plus reader (*BMG LABTECH*). The data was fitted to a segmented linear regression for measuring the compound solubility.

### Cytotoxicity

Cytotoxicity was evaluated in the human neuroblastoma cell line SH-SY5Y (ATCC Number: CRL-2266). Cells were cultured in a mixture of 50% Minimum Essential Medium, 50% Ham’s- F12 medium, 10% fetal bovine serum, 1% L-glutamine, 1% MEM non-essential amino acids and 0.1% of gentamicin. The cells were seeded in 96-well plate at a concentration of 3·10^5^ cells/mL.

After 24 h, the testing compounds were added by triplicate at different concentration up to 100 µM and incubated for further 24 h. Cytotoxicity was sequentially analysed by performing the propidium iodide (PI) fluorescence stain assay and the 3-(4,5-dimethylthiazol-2-yl)-2,5-diphenyl tetrazolium bromide (MTT) colorimetric assay, as described elsewhere (*52*). Briefly, cell death was measured by PI staining. The PI reagent was added to each well and incubated for 1 h, and the fluorescence was measured by Gemini XPS Microplate reader (Millipore) at 530 nm excitation and 645 nm emission, and the relative number of cell death was calculated. Cell metabolic activity was determined by the MTT assay. The MTT reagent was added to each well and incubated for 2 h in the cell incubator, then a lysis buffer was added and further incubated at 37 °C overnight. The colorimetric reaction was measured by Multiskan Spectrum Spectrophotometer (Thermo) at 570 nm and a reference 630 nm wavelength. Experiments were performed three times at different cell passages.

## Supplementary Materials

Fig. S1. Effects of sEH inhibitors on body weight and on water intake in SAMP8 and 5xFAD.

Fig. S2. Significant correlations from the study of SAMP8.

Fig. S3. Significant correlations from the study of 5xFAD.

Fig. S4. Chemical structures of TPPU, UB-EV-52 and AS-2586114, synthesis of the trifluoroacetate salt of AS-2586114 and notation used for the ^1^H and ^13^C NMR assignments.

Table S1. Data Patients.

Table S2. Microsomal stability of UB-EV-52 at human, rat and mice microsomes.

Table S3. Inhibition by UB-EV-52 of recombinant human cytochrome P450 enzymes and of hERG.

Table S4. Partial correlation controlling for group coefficients between selected variables included in the study of SAMP8.

Table S5. Significant correlations from the study of SAMP8.

## Acknowledgments

### Funding

This study was supported by the *Ministerio de Economía, Industria y Competitividad* (Agencia Estatal de Investigación, AEI) and *Fondo Europeo de Desarrollo Regional* (MINECO-FEDER) (Projects SAF2017-82771, SAF2016-77703 and SAF2015-68749) and Generalitat de Catalunya (2017 SGR 106). S.C. and E.P. thank the Universitat de Barcelona and the Institute of Biomedicine of the Universitat de Barcelona (IBUB), respectively, for PhD grants. R.L. and D.P.-I. thank the Spanish Ministerio de Educación, Cultura y Deporte for PhD grants (FPU program). We would like also to thank *Xunta de Galicia (ED431C 2018/21) and Ministry of Economy and Competiveness (Innopharma Project) and European Regional Development Fund (ERDF)*. This study was also supported, in part, by a grant from the National Institute of Environmental Health Sciences (NIEHS) Grant R01 ES002710, and NIEHS Superfund Research Program P42 ES004699. The content is solely the responsibility of the authors and does not necessarily represent the official views of the National Institutes of Health.

### Author contributions

S.V. and M.P. conceived the idea. C.G.-F., S.C., E.P., J.W., R.L., C.E., D. P.-I., J.C., R. C., C.S., M.I.L. and J.B. performed research. C.G.-F., S.V., M.P. and C.G. designed the experiments and analyzed the data. C.M. and B.D.H. provided advice, feedback and the TPPU. C.G.-F., S.V., M.P. and C.G. wrote the.

### Competing interests

C.G.-F., S.C., E.P., R.L., C.E., S.V., M.P. and C.G. are inventors of the Universitat de Barcelona patent applications for the use of sEH inhibitors in the treatment of neurodegenerative diseases (European Patent Application no. 18382266.7 and 18382267.5). S.C., R.L. and S.V. are inventors of the Universitat de Barcelona patent application on sEH inhibitors (WO2017017048). C.M. and B.D.H. are inventors of the University of California patents on sEH inhibitors licensed to EicOsis (WO2012054093). None of the other authors has any disclosures to declare.

### Data and materials availability

UB-EV-52 must be obtained through an MTA.

