## Supplementary data for "Pharmacological inhibition of soluble epoxide hydrolase as a new therapy for Alzheimer’s Disease"

**Supplementary Materials:**

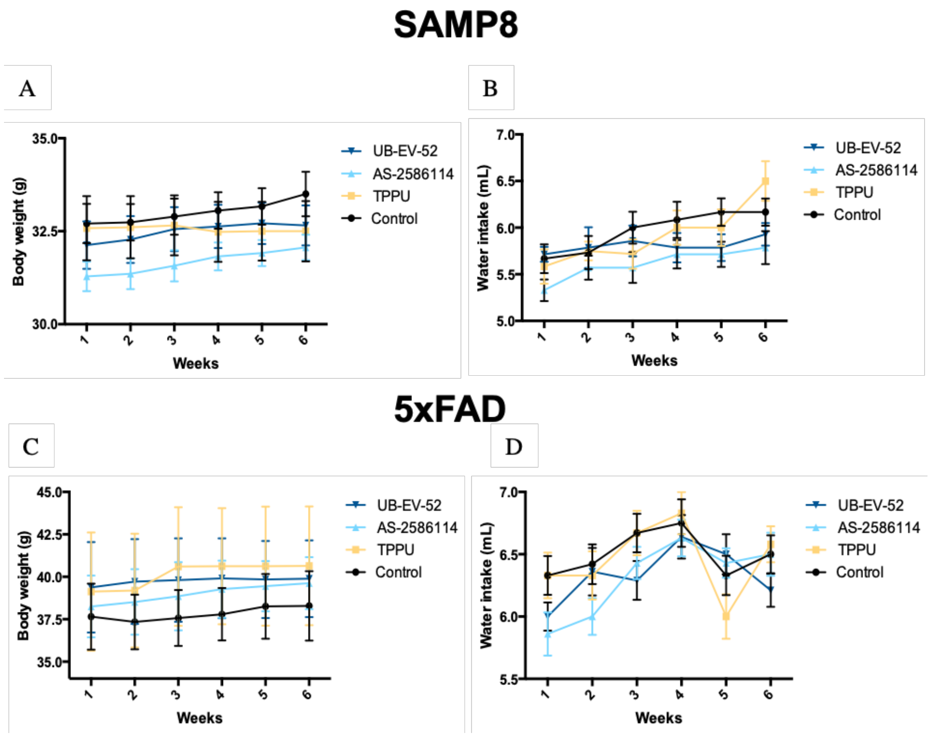


**Fig. S1. Effects of sEH inhibitors on body weight and on water intake in SAMP8 and 5xFAD.** **a.** Body weight in SAMP8; body weight was measured every week during the treatment (n = 12-14 per group). **b.** Water intake in SAMP8; mice were administered by drinking water. **c.** Body weight in SAMP8; body weight was measured every week during the treatment (n = 12-14 per group). **d.** Water intake in 5xFAD; mice were administered by drinking water. The water intake amount was recorded every week throughout the treatment (n = 12-14 per group). Results are expressed as a MEAN ± SEM and were not significantly different from the control group. Groups were compared by Student t-test or by One-Way ANOVA and post-hoc Dunnett's, n = 12-14 per group, (*Significant at $p<0.05$).

### SAMP8

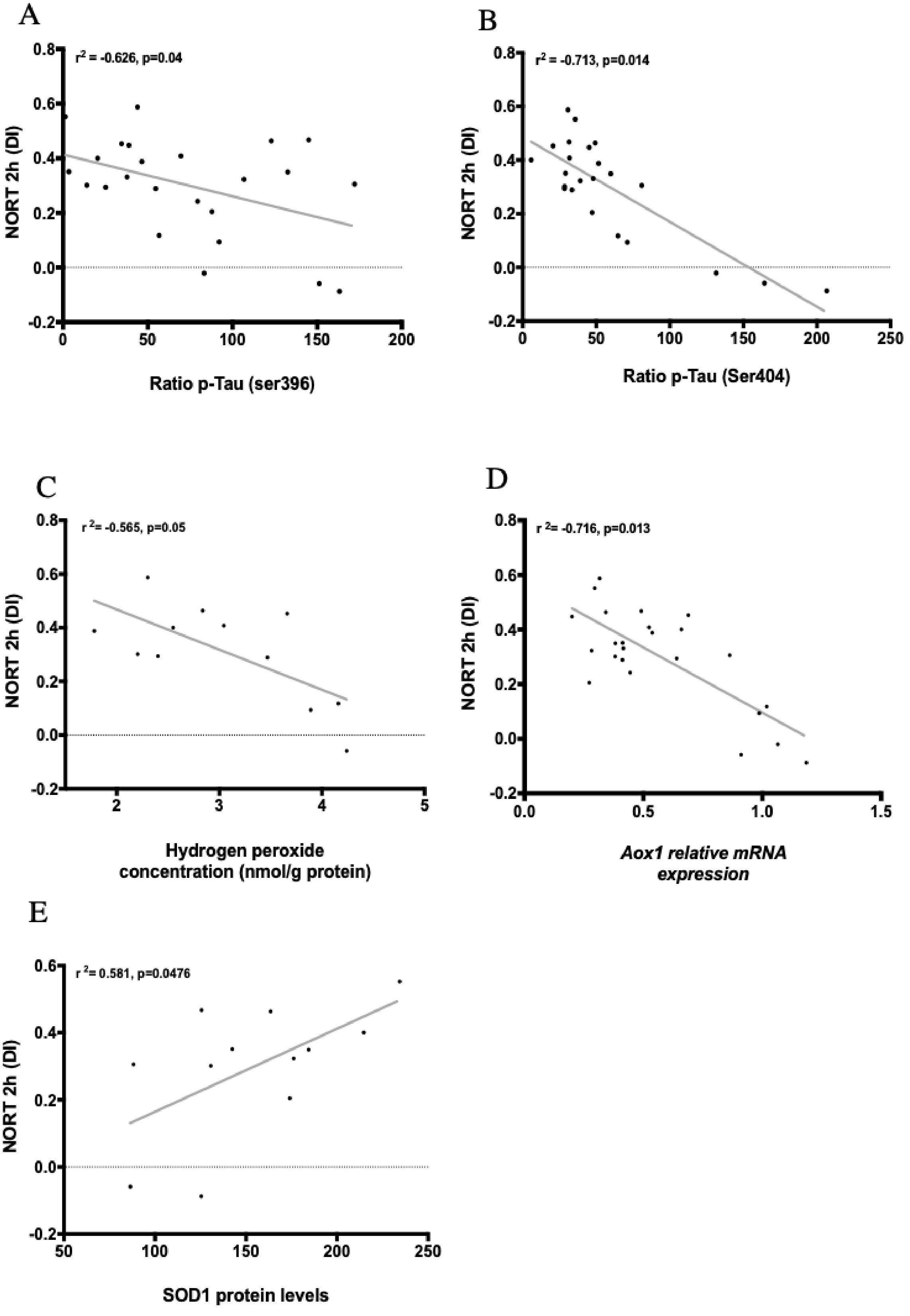

A
$r^2$ = -0.626, p=0.04
NORT 2h (DI)
Ratio p-Tau (ser396)
B
$r^2$ = -0.713, p=0.014
NORT 2h (DI)
Ratio p-Tau (Ser404)
C
$r^2$= -0.565, p=0.05
NORT 2h (DI)
Hydrogen peroxide concentration (nmol/g protein)
D
$r^2$= -0.716, p=0.013
NORT 2h (DI)
*Aox1 relative mRNA expression*
E
$r^2$= 0.581, p=0.0476
NORT 2h (DI)
SOD1 protein levels

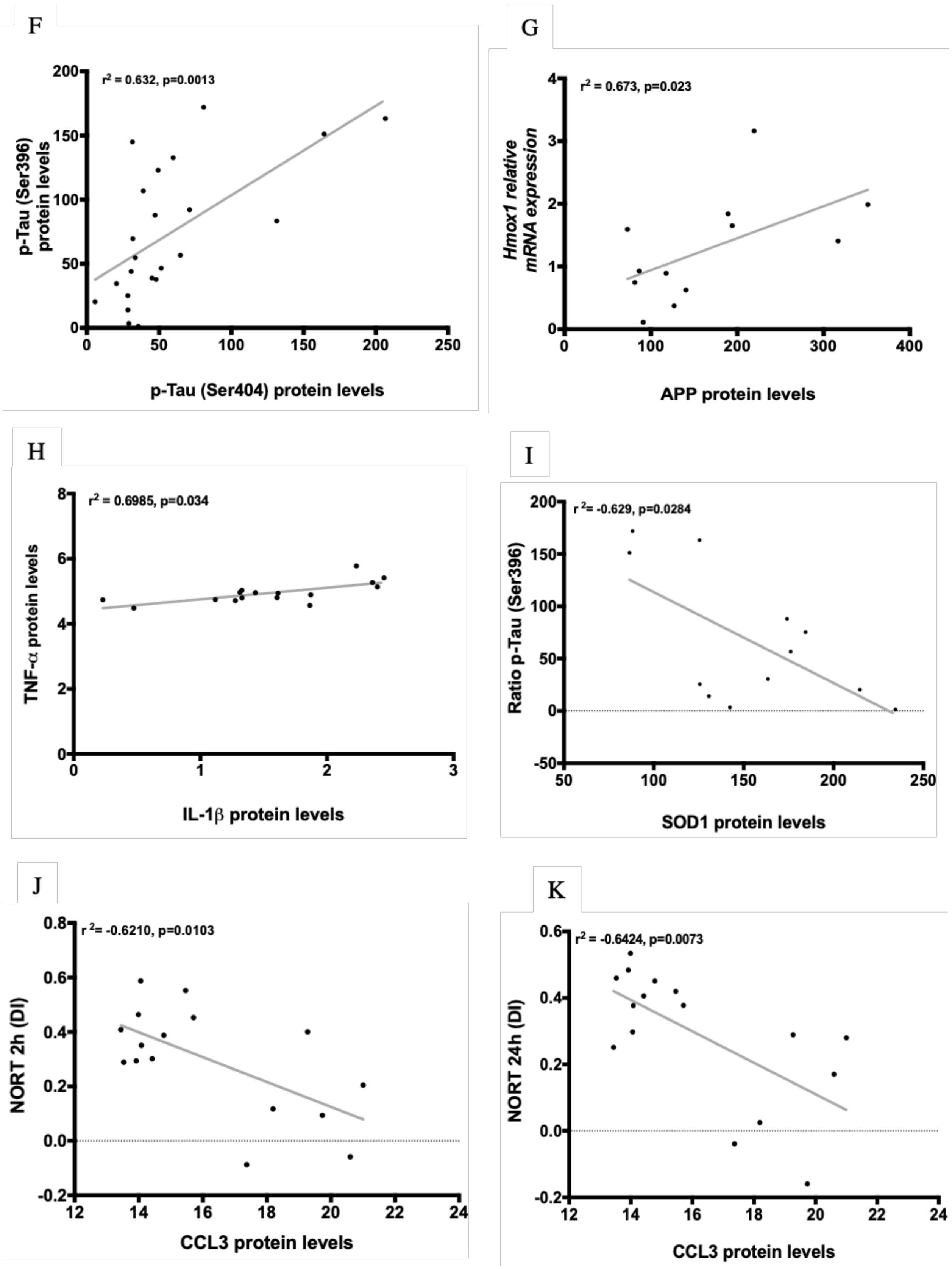
F
r2 = 0.632, p=0.0013
p-Tau (Ser396) protein levels
p-Tau (Ser404) protein levels
G
r2 = 0.673, p=0.023
Hmox1 relative mRNA expression
APP protein levels
H
r2 = 0.6985, p=0.034
TNF-α protein levels
IL-1β protein levels
I
r 2= -0.629, p=0.0284
Ratio p-Tau (Ser396)
SOD1 protein levels
J
r 2= -0.6210, p=0.0103
NORT 2h (DI)
CCL3 protein levels
K
r2= -0.6424, p=0.0073
NORT 24h (DI)
CCL3 protein levels

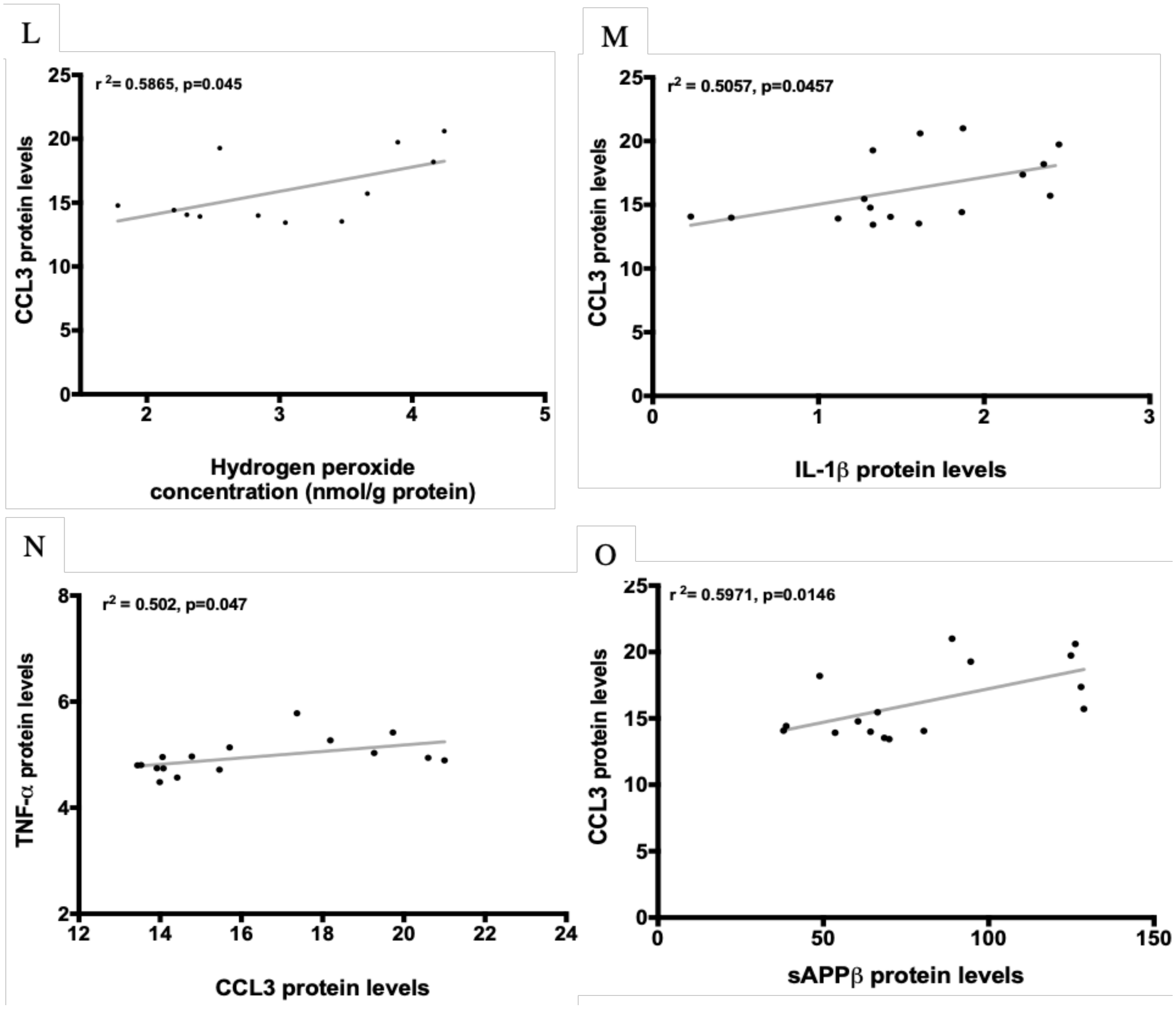


**Figure S2. Significant correlations from the study of SAMP8.** Correlations between Ratio p-Tau Ser396 **(a)**, Ratio p-Tau Ser404 **(b)**, Hydrogen peroxide concentration **(c),** *Aox1* gene expression **(d),** and SOD1 protein levels **(e)** with NORT 2h (DI). Correlation between Ratio p-Tau Ser404 **(f)** with Ratio p-Tau Ser396. Correlation between APP protein levels **(g)** with *Hmox1* gene expression. Correlation between IL-1β **(h)** with TNF-α protein levels. Correlation between SOD1 **(i)** with Ratio p-Tau Ser396. Correlations between NORT 2h (DI) **(j)**, NORT 24h (DI) **(k)**, Hydrogen peroxide concentration **(l)**, IL-1β **(m)**, TNF-α **(n)**, and sAPPβ **(o)** with CCL3 protein levels. Partial correlation was performed between protein levels of Tau pathology, oxidative stress, inflammatory and APP processing markers in the hippocampus and behavioural test parameters of all groups of SAMP8 (n = 14). $R^2$ and p-values are indicated on graphs, (*Significant at $p<0.05$).

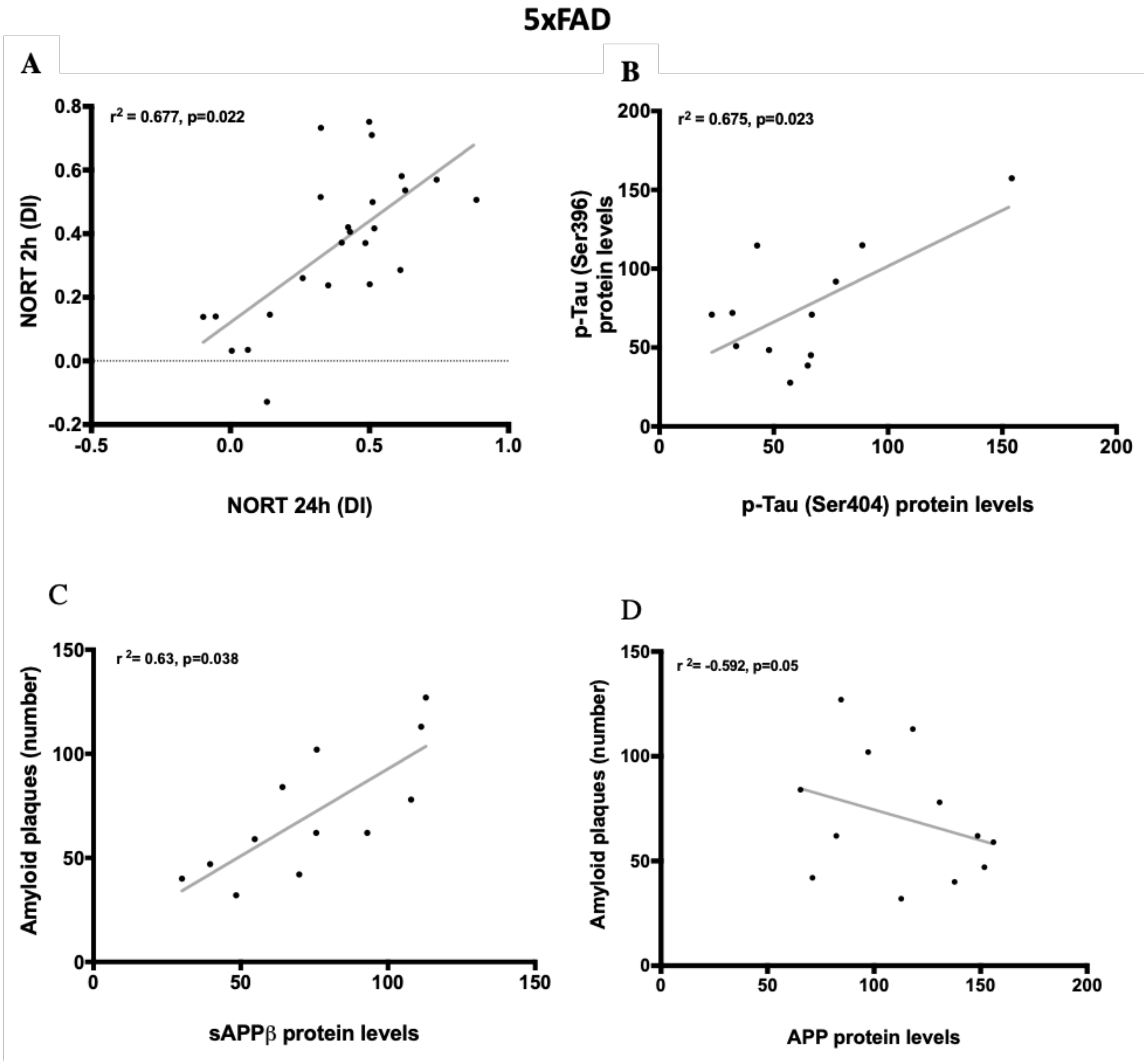


**Fig. S3. Significant correlations from the study of 5xFAD.** Correlation between NORT 24h (DI) **(A)** with NORT 2h (DI). Correlation between Ratio p-Tau Ser396 **(B)** with Ratio p-Tau Ser404. Correlations between sAPPβ **(C),** and APP protein levels **(D)** with number of amyloid plaques. Partial correlation was performed between protein levels of Tau pathology and APP processing markers in the hippocampus and behavioural test parameters of all groups of 5xFAD (n = 12-14). $R^2$ and p-values were indicated on graphs. (*Significant at $p<0.05$).

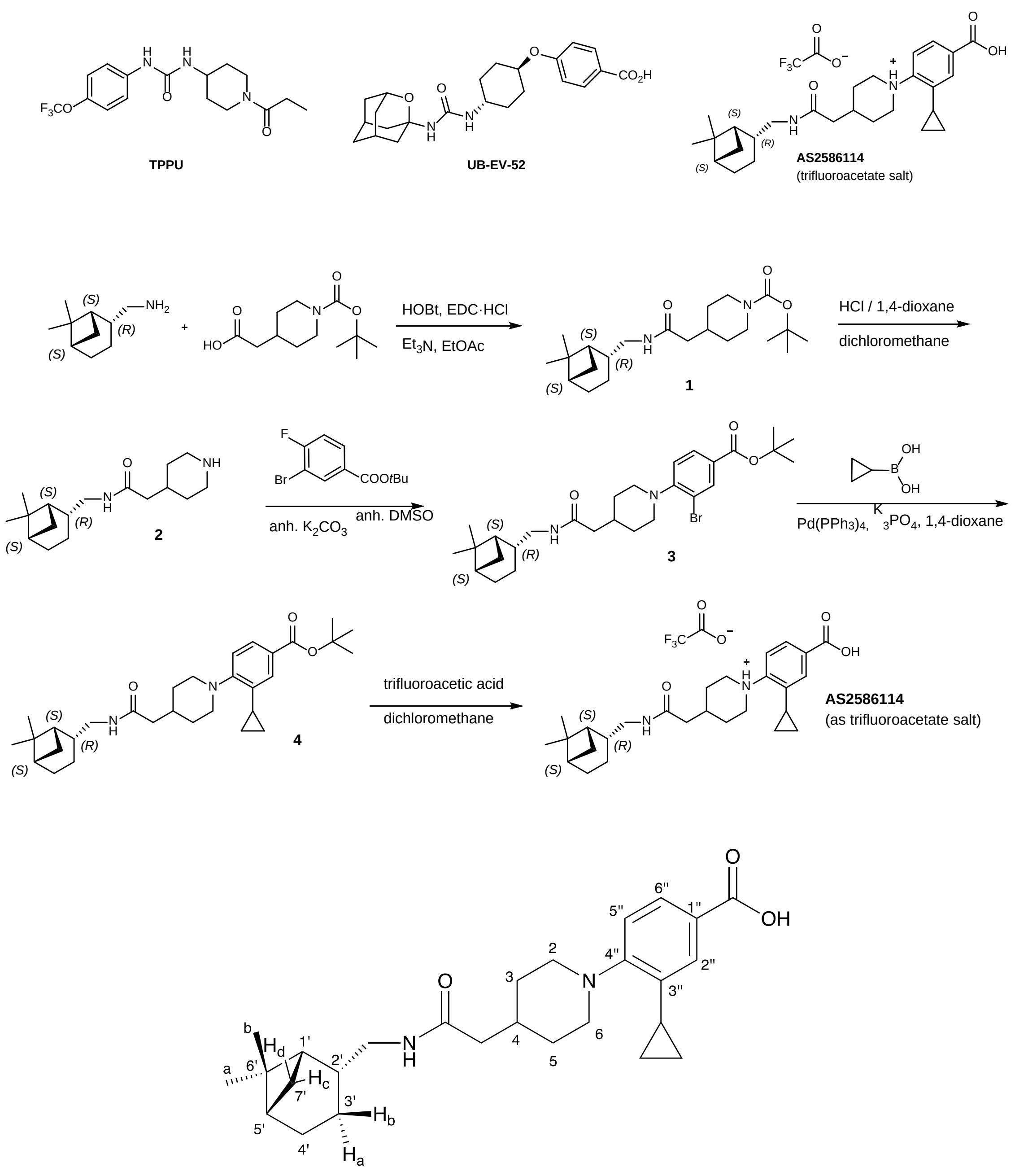


**Fig. S4. Chemical structures of TPPU, UB-EV-52 and AS-2586114, synthesis of the trifluoroacetate salt of AS-2586114 and notation used for the $^{1}$H and $^{13}$C NMR assignments**.

| GENDER | AGE | POSTMORTEM DELAY | DIAGNOSIS |
|---|---|---|---|
| Male | 85 | 5 h 45 min | Control |
| Male | 78 | 2 h 15 min | Control |
| Male | 79 | 7 h | Control |
| Male | 93 | 7 h 20 min | AD BRAAK STAGE V/C |
| Female | 82 | 1 h 45 min | AD BRAAK STAGE V/B |
| Female | 72 | 9 h 30 min | AD BRAAK STAGE V/B |
| Female | 75 | 4 h 15 min | AD BRAAK STAGE V/B |

**Table S1. Data Patients.**

| Human | | | Rat | | | Mice | | |
|---|---|---|---|---|---|---|---|---|
| % remanent (60 min) | $t_{1/2}$ (min) | Clint (µL/min*mg prot) | % remanent (60 min) | $t_{1/2}$ (min) | Clint (µL/min*mg prot) | % remanent (60 min) | $t_{1/2}$ (min) | Clint (µL/min*mg prot) |
| 50 | 57 | 8.1 | 92 | 496 | 1.9 | 90 | 396 | 2.3 |

**Table S2. Microsomal stability of UB-EV-52 at human, rat and mice microsomes**

| CYP1A2 | CYP2C9 | CYP2C19 | CYP3A4 (BFC) | CYP3A4 (DBF) | CYP2D6 | hERG |
|---|---|---|---|---|---|---|
| 1 ± 1 | 3 ± 1 | 4 ± 1 | 1 ± 1 | 29 ± 1 | 2 ± 1 | 14 ± 4 |

**Table S3. Inhibition by UB-EV-52 of recombinant human cytochrome P450 enzymes (%inhib at 10 µM) and of hERG channel (%inhib at 10 µM).**

| | **DI 2h** | **DI 24h** | **p-Tau Ser396** | **p-Tau Ser404** | **ROS** | ***Aox1*** | ***Hmox1*** | **SOD1** | ***Il-1β*** | ***Tnf-α*** | ***Ccl3*** | **IL-1β** | **TNF-α** | **CCL3** | **ATF-6** | **IRE1α** | **XBP1** | **APP** | **sAPPα** | **sAPPβ** | **β-CTF** |
|---|---|---|---|---|---|---|---|---|---|---|---|---|---|---|---|---|---|---|---|---|---|
| **DI 2h** | 1 | 0,417 | **-0,626**$^{*}$ | **-0,713**$^{*}$ | **-0,565**$^{*}$ | **-0,716**$^{*}$ | 0,091 | **0,581**$^{*}$ | -0,254 | -0,377 | -0,317 | -0,24 | -0,034 | **-0,621**$^{*}$ | -0,350 | -0,276 | -0,401 | -0,126 | 0,046 | 0,09 | 0,076 |
| **DI 24h** | | 1 | -0,19 | -0,043 | 0,009 | -0,53 | 0,141 | 0,306 | -0,334 | -0,39 | -0,268 | -0,482 | -0,535 | **-0,642**$^{**}$ | -0,432 | -0,201 | -0,391 | -0,284 | -0,043 | 0,198 | 0,316 |
| **p-Tau Ser396** | | | 1 | **0,632**$^{**}$ | 0,392 | 0,248 | -0,59 | **-0,629**$^{*}$ | -0,108 | -0,018 | -0,066 | -0,127 | -0,147 | 0,434 | -0,211 | -0,104 | -0,259 | -0,206 | -0,289 | 0,273 | 0,069 |
| **p-Tau Ser404** | | | | 1 | 0,34 | 0,296 | -0,363 | -0,310 | 0,039 | 0,179 | 0,098 | -0,196 | -0,304 | 0,428 | -0,294 | -0,298 | -0,375 | -0,1 | -0,23 | 0,088 | 0,111 |
| **ROS** | | | | | 1 | 0,369 | -0,411 | 0,005 | 0,037 | -0,015 | 0,015 | 0,269 | -0,147 | **0,586**$^{*}$ | -0,462 | -0,262 | -0,346 | -0,374 | -0,173 | 0,025 | 0,045 |
| ***Aox1*** | | | | | | 1 | 0,227 | -0,482 | -0,041 | 0,163 | 0,02 | 0,417 | 0,538 | 0,488 | 0,025 | 0,034 | 0,215 | 0,286 | 0,199 | -0,008 | -0,055 |
| ***Hmox1*** | | | | | | | 1 | 0,301 | 0,202 | 0,307 | 0,325 | 0,155 | 0,206 | 0,269 | 0,119 | 0,126 | 0,324 | **0,673**$^{*}$ | 0,565 | -0,01 | -0,126 |
| **SOD1** | | | | | | | | 1 | 0,323 | -0,299 | -0,186 | -0,199 | -0,211 | -0,094 | 0,092 | 0,024 | 0,210 | 0,001 | -0,278 | -0,375 | -0,437 |
| ***Il-1β*** | | | | | | | | | 1 | 0,939 | 0,976 | 0,44 | -0,218 | -0,402 | 0,203 | 0,241 | 0,294 | 0,467 | 0,306 | -0,249 | -0,381 |
| ***Tnf-α*** | | | | | | | | | | 1 | 0,954 | 0,461 | -0,122 | 0,078 | 0,106 | 0,218 | 0,319 | 0,537 | 0,42 | -0,286 | -0,196 |
| ***Ccl3*** | | | | | | | | | | | 1 | 0,415 | -0,228 | -0,05 | -0,015 | -0,212 | -0,323 | 0,546 | 0,42 | -0,152 | -0,359 |
| **IL-1β** | | | | | | | | | | | | 1 | **0,698**$^{*}$ | **0,505**$^{*}$ | 0,289 | 0,123 | 0,242 | 0,571 | 0,439 | 0,034 | -0,054 |
| **TNF-α** | | | | | | | | | | | | | **1** | **0,502**$^{*}$ | 0,367 | 0,254 | 0,395 | 0,497 | 0,214 | 0,132 | -0,008 |
| **CCL3** | | | | | | | | | | | | | | **1** | 0,253 | 0,311 | 0,258 | -0,175 | -0,062 | **0,597**$^{*}$ | 0,495 |
| **ATF-6** | | | | | | | | | | | | | | | **1** | 0,321 | 0,426 | 0,224 | -0,235 | 0,346 | 0,034 |
| **IRE1α** | | | | | | | | | | | | | | | | **1** | 0,214 | 0,045 | 0,042 | 0,216 | -0,021 |
| **XBP1** | | | | | | | | | | | | | | | | | **1** | 0,324 | -0,367 | -0,379 | 0,245 |
| **APP** | | | | | | | | | | | | | | | | | | 1 | 0,549 | 0,234 | -0,37 |
| **sAPPα** | | | | | | | | | | | | | | | | | | | 1 | 0,392 | -0,042 |
| **sAPPβ** | | | | | | | | | | | | | | | | | | | | **1** | -0,349 |
| **β-CTF** | | | | | | | | | | | | | | | | | | | | | 1 |

**Table S4. Partial correlation controlling for group coefficients between selected variables included in the study of SAMP8**. The values used to calculate Partial correlation controlling for group coefficients were behavioral parameter from Figure 5, gene expression (shown in Figures 3 and 4) and protein levels (shown in Figures 3 and 4). Correlation (2-tailed) is significant *p<0.05; **p<0.01; (-) Negative covariation of two variables.

| | DI 2h | DI 24h | p-Tau Ser396 | p-Tau Ser404 | APP | sAPPα | sAPPβ | β-CTF | Amyloid plaques |
|---|---|---|---|---|---|---|---|---|---|
| **DI 2h** | 1 | **-0,677*** | -0,146 | 0,038 | -0,316 | 0,477 | 0,074 | -0,253 | 0,038 |
| **DI 24** | | 1 | -0,3 | -0,19 | -0,119 | 0,439 | 0,038 | -0,311 | -0,329 |
| **p-Tau Ser396** | | | 1 | **0,675*** | 0,01 | -0,177 | 0,002 | 0,079 | 0,363 |
| **p-Tau Ser404** | | | | 1 | -0,01 | 0,334 | 0,195 | 0,014 | 0,12 |
| **APP** | | | | | 1 | -0,347 | -0,329 | -0,266 | **-0,592*** |
| **sAPPα** | | | | | | 1 | 0,166 | 0,015 | -0,015 |
| **sAPPβ** | | | | | | | 1 | -0,124 | **0,638*** |
| **β-CTF** | | | | | | | | 1 | 0,181 |
| **Amyloid plaques** | | | | | | | | | 1 |

**Table S5. Partial correlation controlling for group coefficients between selected variables included in the study of 5xFAD**. The values used to calculate Partial correlation controlling for group coefficients were behavioral parameter from Figure 4, protein levels (shown in Figure 3) and amyloid plaques (shown in Figure 3). Correlation (2-tailed) is significant *p<0.05; (-) Negative covariation of two variables.
